# Correlated cryogenic fluorescence microscopy and electron cryotomography shows that exogenous TRIM5α can form hexagonal lattices or autophagy aggregates in vivo

**DOI:** 10.1101/835322

**Authors:** Stephen D. Carter, João I. Mamede, Thomas J. Hope, Grant J. Jensen

## Abstract

Members of the TRIM protein family have been shown to gather into structures in both the nucleus and cytoplasm. One TRIM protein family member, TRIM5α, has been shown to form cytoplasmic bodies involved in restricting retroviruses such as HIV-1. Here we applied cryogenic correlated light and electron microscopy (cryo-CLEM) to intact mammalian cells expressing YFP-rhTRIM5α and found hexagonal nets were present whose arm-lengths were similar to those of the hexagonal nets formed by purified TRIM5α in-vitro. We also observed YFP-rhTRIM5α within a diversity of structures with characteristics expected for organelles involved in different stages of macroautophagy, including disorganized protein aggregations (sequestosomes), sequestosomes flanked by flat double-membraned vesicles (sequestosome:phagophore complexes), sequestosomes within a double-membraned vesicle (autophagosomes), and sequestosomes within multi-vesicular autophagic vacuoles (autolysosomes or amphisomes). Vaults were also seen in these structures, consistent with their role in autophagy. Our data (i) support recent reports that TRIM5α can form both well-organized signaling complexes and non-signaling aggregates, (ii) offer the first images of the macroautophagy pathway in a near-native state, and (iii) reveal that vaults arrive early in macroautophagy.

## Introduction

Members of the tripartite motif (TRIM) family of proteins have been shown to localize to isolated subcellular compartments ^1^. One of the most studied TRIM compartments is the TRIM5α cytoplasmic body ^2^. Similar to other members of the TRIM family ^1^, TRIM5 proteins comprise a RING E3 ubiquitin-ligase domain, a B-box 2 self-assembly domain, and an antiparallel dimeric coiled-coil. Li et al. showed previously that the coiled-coil and B-box 2 domains of TRIM5α facilitate oligomerization into flat hexagonal lattices *in vitro* ^3–6^. TRIM5 proteins also contain one of two different C-terminal viral recognition domains, a B30.2/SPRY domain in TRIM5α or a cyclophilin A (CypA) domain in TRIMCyp ^7–10^. One of the most notable features of TRIM5 proteins is their ability to block retroviral infections by binding viral capsids via their B30.2/SPRY or CypA domains ^2,10^, but how this leads to restriction remains unclear.

Campbell et al. reported previously that fluorescently-tagged human TRIM5α (huTRIM5α) and rhesus TRIM5α (rhTRIM5α) form highly dynamic fluorescent bodies of various sizes inside cells ^11,12^, which interact with cytoplasmic HIV-1 viral complexes ^12^. Previous FIB-SEM studies revealed that fixed and stained YFP-tagged rhesus TRIM5α (YFP-rhTRIM5α) bodies appear as large aggregates inside the cytoplasm ^13^, but no further detail was discernable. Towards elucidating the ultrastructure of YFP-rhTRIM5α bodies to higher resolution and in a more native state, here we sought to image them using cryogenic correlated light and electron microscopy (cryo-CLEM) ^14^.

We find that YFP-rhTRIM5α can form extended hexagonal nets inside cells as well as different structures whose features match those expected for different stages of macroautophagy, including sequestosomes^15^, phagophores, autophagosomes and autophagic vacuoles. Macroautophagy is one form of autophagy in which large structures such as mitochondria, misfolded and aggregated proteins, viruses, or other organelles or cellular pathogens are degraded ^16,17^. The process begins with sequestration of the substrates within the cytoplasm. Then a cup-shaped, flattened vesicle called a phagophore grows around the sequestered material until it fuses with itself, enclosing the substrates within a double-membraned autophagosome.

Autophagosomes then travel long distances along microtubules towards the microtubule-organizing center until they encounter and fuse with either an endosome or a lysosome to form an amphisome or auto-lysosome, respectively. Membrane fusion of an amphisome or autophagosome with a lysosome can also produce an auto-lysosome. Another structure known to be involved in autophagy are vaults. Vaults are 70-nm by 30-nm, roughly-ellipsoidal, megadalton ribonucleoprotein complexes comprising the major vault protein (MVP), telomerase-associated protein-1 (TEP1), and the vault poly (ADP-ribose) polymerase. Vaults reside in the cytoplasm ^44^ and nucleus and are known to house non-coding vault RNAs (vtRNAs) important in the regulation of autophagy ^45^.

As a result of finding that YFP-rhTRIM5α bodies are associated with macroautophagy, our cryotomograms provide 3D images of all these structures in their native state to macromolecular (~4 nm) resolution, and reveal that vaults arrive early in the process.

## Results

HeLa cells either stably overexpressing YFP-rhTRIM5α or not (wildtype controls) were grown on EM grids for 12 hours and then plunge-frozen in liquid ethane ^18^. Because the proteasome inhibitor MG-132 generates large YFP-rhTRIM5α fluorescent bodies and was shown to have little effect on steady state protein levels of YFP-rhTRIM5α (compared to untreated) ^12,19^, cells treated with MG-132 were also imaged (see Table S2 for details on which tomograms came from which cells). Phase contrast and fluorescence images were then recorded of the frozen cells using a light microscope equipped with a cryo-stage and a long-working-distance air-objective lens ^14^. Grids with YFP-rhTRIM5α fluorescent bodies present in cell peripheries suitably thin for transmission electron microscopy were then transferred into a cryo-electron microscope. These same cell regions were then re-located within the cryo-electron microscope and full tilt-series were recorded. Three-dimensional reconstructions (cryotomograms) were calculated. 500 nm blue fluorospheres added to the samples before freezing were used as fiducial markers to precisely superimpose the centroids of the fluorescent puncta within 80 nm accuracy onto the cryotomograms, using the same image correlation procedures described previously (Figure S1) ^20^.

Thirty-five cryotomograms of YFP-TRIM5α fluorescent bodies located in the cell periphery of 20 cells and 51 control cryotomograms of HeLa cells not expressing YFP-rhTRIM5α were analyzed in total. As we inspected the cryotomograms of YFP-TRIM5α bodies, and compared them to the controls, we recognized several structures frequently co-localizing with YFP-TRIM5α including vaults, dense protein aggregates, flattened vesicles, double-membraned vesicles, and nested vesicles. Because all these structures are expected for the autophagy pathway, we concluded that exogenous YFP-TRIM5α was being degraded through autophagy. This allowed us to begin to place the cryotomograms in sequence within the autophagy process based on both which features accompanied which others (protein aggregates were sometimes present without flattened vesicles, but not vice-versa) and existing knowledge, including for instance that phagophores (the flattened vesicles) are known to mature into double-membraned autophagosomes. This analysis revealed distinct stages of autophagy and their order: colocalization of substrate, arrival of vaults, formation of dense substrate aggregates (sequestosomes), creation of phagophores, phagophore closure around substrate to produce autophagosomes, and finally autophagosome fusion with either endosomes or lysosomes to produce an autophagic vacuole.

In the figures here, we present example cryotomograms representing each of the different stages of autophagy in their deduced sequence. For clarity and simplicity, each such figure shows one cryotomogram, with multiple panels to highlight different features. Figures 1 and S2 show vaults; Figures 2 and S3-5 show pre-phagophore sequestosomes; Figures 3, 4, and S6 show sequestosome:phagophore complexes; Figures 6 and S8-9 show autophagosomes; and Figures 7 and S10 show autophagic vacuoles. Again for clarity and simplicity, these figure panels follow a consistent pattern: the first panel shows an overlay of the yellow-channel fluorescence image (revealing the location of YFP-rhTRIM5α) on an xy slice of the cryotomogram, the second panel shows the same slice of the cryotomogram without the fluorescence, the third panel shows a color-coded segmentation of the 3D cellular ultrastructure as revealed by the cryotomograms, and the fourth panel shows an enlarged view of key structures of interest, sometimes rotated to a more useful perspective. In some cases, additional panels or figures are shown to highlight other important findings, and three movies showing the fluorescence images, correlation process, cryotomograms, and 3D segmentations from various angles of the volumes shown in Figures 2, 4 and S6 are provided as Movies 1, 2 and 3. The pre-figure illustrates the color code used in the segmentations to mark the different organelles.

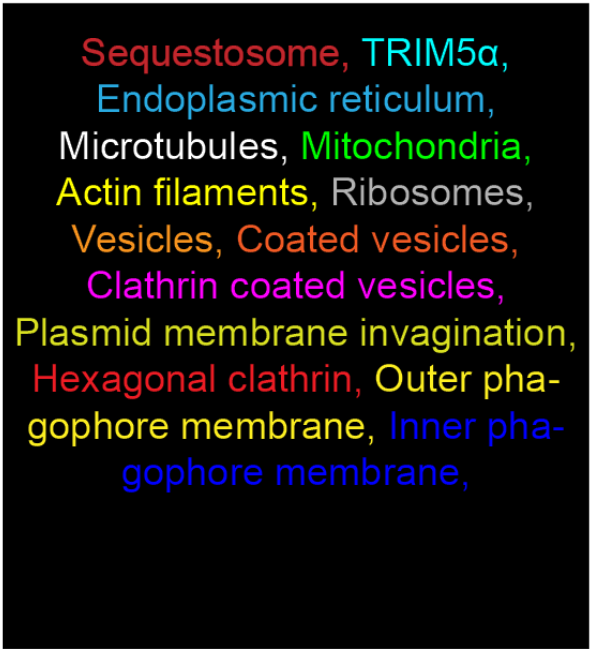

**Figure 1.**
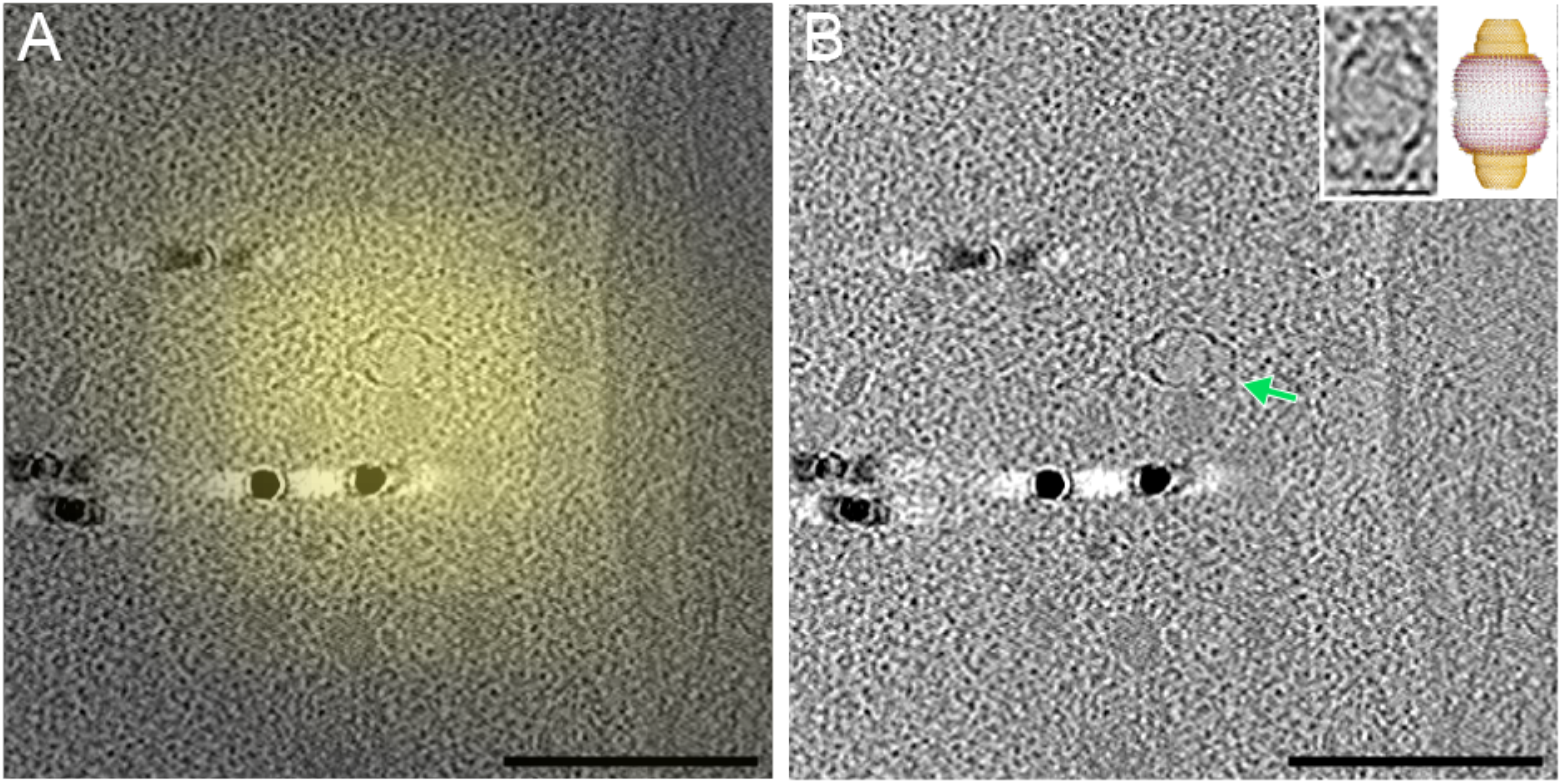
Here and all other main figures below A, B, C and D panels are arranged as follows (A) Deconvolved epifluorescent images overlaid onto a cryotomogram slice. (B) Cryotomographic slice without fluorescence. (C) Segmentation of the 3D ultrastructure inside the cell. (D) Middle right panel shows a zoomed view of the sequestosome, hexagonal clathrin and TRIM5α nets in (B) and (C). Cellular structures are labelled as followed ER (endoplasmic reticulum), M (mitochondrion), AV (autophagic vacuole), MVB (multilamella fluorescent bodies), V (vesicle) MT (microtubule), P (phagophore), A (actin filaments), F (filapodia), and AP (autophagosome). **Cryo-CLEM reveals YFP-TRIM5α fluorescent bodies localize near vault complexes.** The green arrow highlights a vault complex. Scale bars = 150 nm. Inset shows an enlarged view of a vault, alongside the crystal structure (PDB:4V60) of the vault complex. Scale bar = 30 nm.

**Figure 2.**
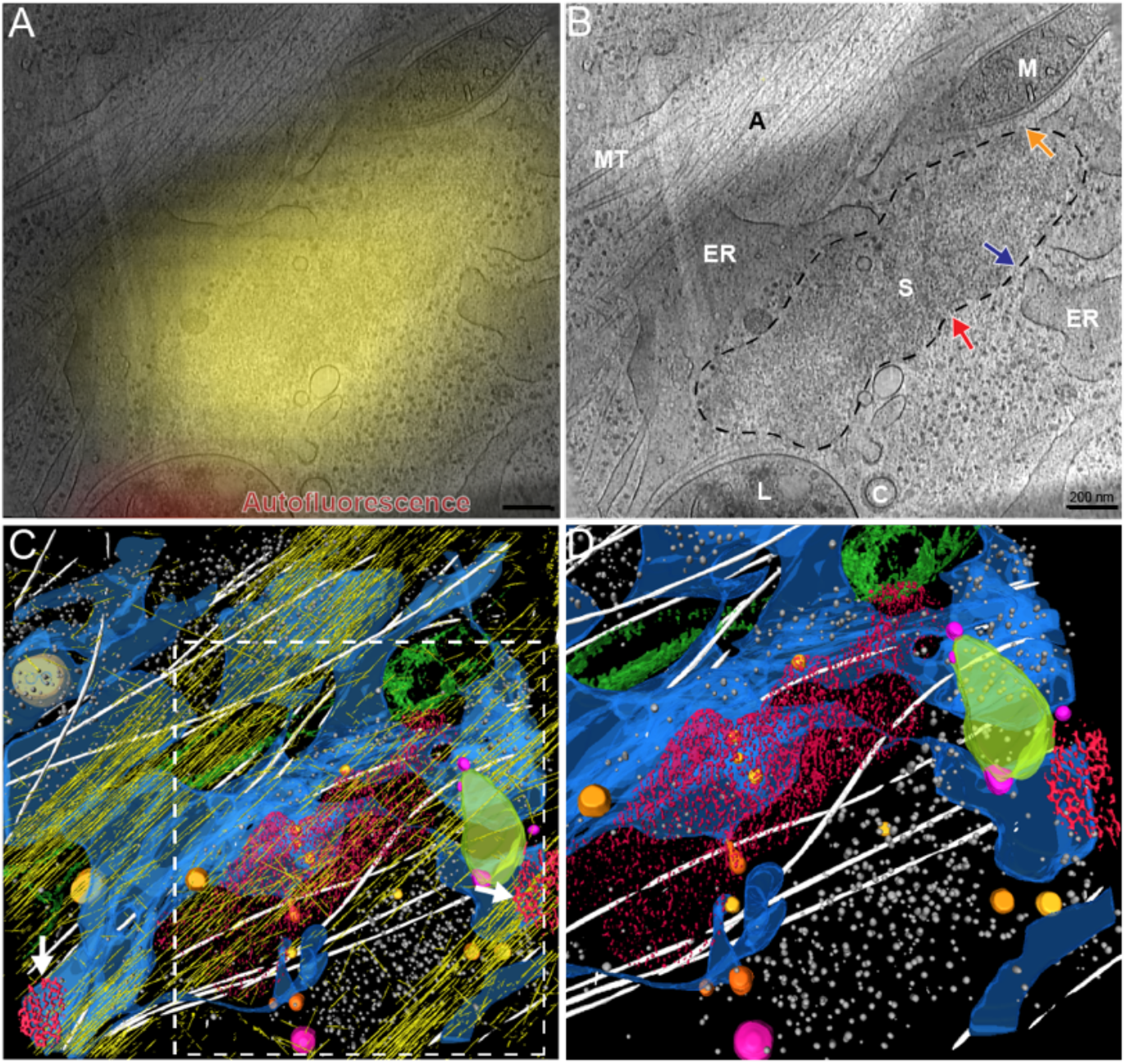
Sequestosomes represent YFP-rhTRIM5α fluorescent bodies in MG-132 treated HeLa cells. **(B)** Same cryotomogram slice as A, with the border of the sequestosome highlighted with a black dashed line. The red arrow highlights a cytoplasm:sequestosome boundary, the orange arrow highlights a mitochondrion:sequestosome boundary, and the blue arrow highlights an ER:sequestosome in close proximity. A and B Scale bars = 200 nm. **(C)** The two white arrows point to hexagonal clathrin at the plasma membrane. **(D)** Enlarged view of the ER, sequestosome, and hexagonal clathrin, with the actin filaments omitted, as highlighted by the white dashed box in **(C)**.

**Figure 3.**
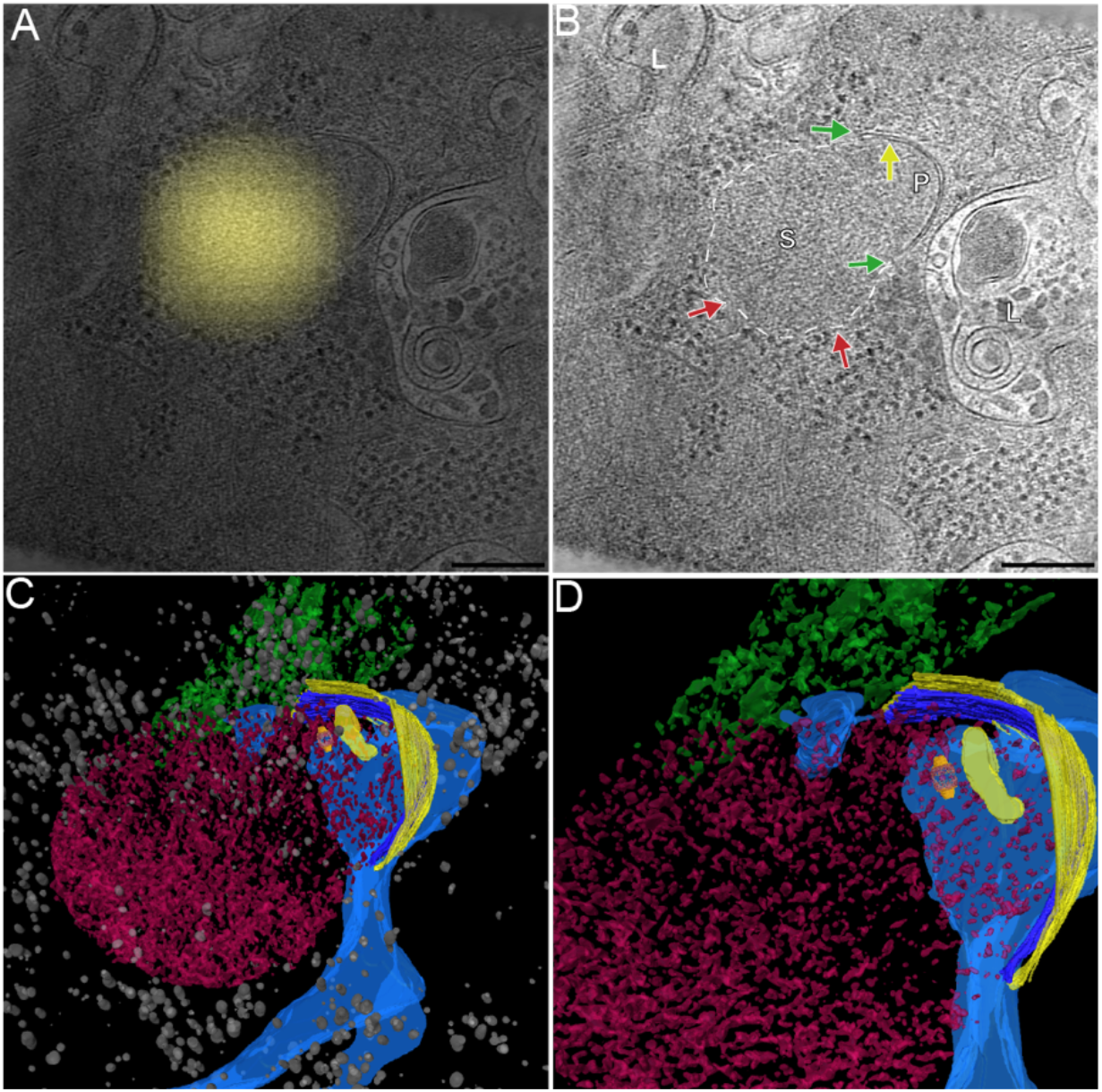
Cryo-CLEM reveals YFP-rhTRIM5α fluorescent bodies (untreated), localized to a cytosolic sequestosome:phagophore complex in close association with the ER. **(B)** The boarder of the sequestosome is highlighted with a white dashed line. The yellow arrow highlights a sequestosome:phagophore in close proximity, the red arrow highlight a cytoplasm:sequestosome in close proximity, and the green arrows highlight the extreme ends of the phagophore membrane. Scale bars = 200 nm. **(D)** Enlarged view of the sequestosome:phagophore complex shown in **(B)** and **(C)**.

**Figure 4.**
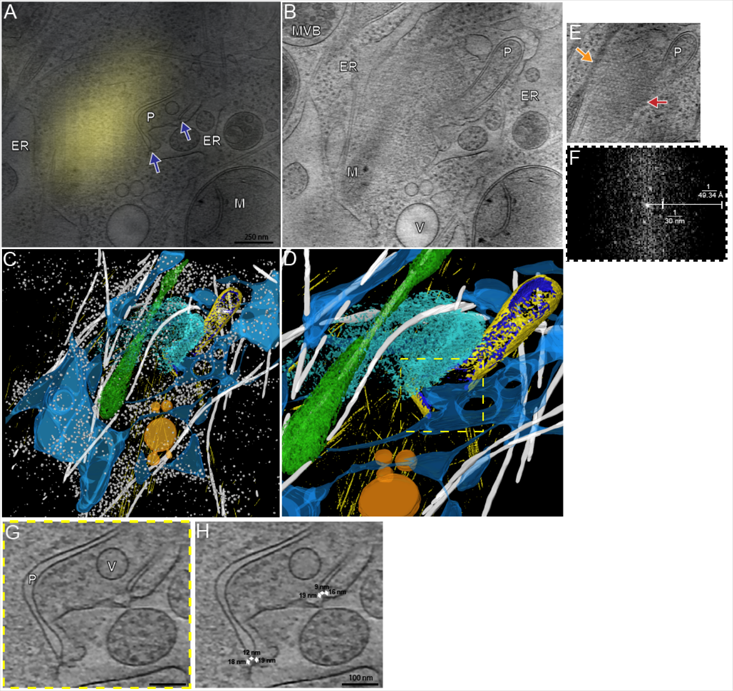
Phagophores in association with a TRIM5α hexagonal lattice represent YFP-rhTRIM5α fluorescent bodies. **(A)** Cryotomographic slice displaying the base of the phagophore. The blue arrows highlight the close proximity between the extreme ends of the phagophore and ER. **(B)** A different cryotomographic slice displaying the tip of the phagophore. A and B Scale bars = 250 nm. **(D)** Enlarged view of the TRIM5α hexagonal lattice shown in **(B)** and **(C)**. **(E)** Cryotomographic slice showing TRIM5α hexagonal nets found located in the cytoplasm. The red arrow highlights a cytoplasm:TRIM5α hexagonal lattice in close proximity, the orange arrow highlight mitochondrion:TRIM5α hexagonal lattice in close proximity. Scale bar = 100 nm. **(F)** Displays a Fourier transform of the cytosolic hexagonal TRIM5α. **(G)** An enlarged image of the extreme ends of the phagophore tethered to the ER membrane via TRIM5α arm-like protein densities, as highlighted by the yellow dashed box in **(D)**. **(H)** The same image as in **(G)**, but this time with the distances of the arm-like densities in nm. Scale bar = 100 nm.

### Stage 1: Colocalization of substrate

In 5 of the cryotomograms of YFP-TRIM5α fluorescent bodies, none of the above-mentioned hallmarks of autophagy were found, and the scene was just typical cytoplasm (as in the controls). Our interpretation is that autophagy begins by substrate colocalization that is sufficient to form a fluorescent punctum, but without structural features recognizable by cryo-ET.

### Stage 2: Vault complexes

Vaults, with their unique and easily-recognizable morphology, were seen in 7 of the 35 total cryotomograms. Because no vaults were seen in the control cryotomograms of HeLa cells not expressing YFP-rhTRIM5α, we concluded that vaults are hallmarks of the autophagy pathway, rather than just ubiquitously present in the cytoplasm. Vaults were seen in cryotomograms alone (without any other hallmarks of autophagy visible) and at all other stages (with sequestosomes, phagophores, autophagosomes, and within autophagic vacuoles). Because vaults were seen alone in two cases (shown in Fig. 1 and S2), but also in all other stages of autophagy, we interpret these two cryotomograms as early stages of autophagy, and conclude that vaults are recruited before visible sequestosomes form (see below).

### Stage 3: Pre-phagophore sequestosomes

Ten of the 28 YFP-rhTRIM5α fluorescent bodies co-localized with disorganized protein aggregates in the midst of unstructured cytoplasm that were recognizable both from their unique texture of packed dot-like densities and exclusion of organelles and vesicles. We applied a convolution neural network to objectively identify and annotate ribosomes and microtubules ^21^, and found that they were excluded from the aggregates. (Figures. 2B red arrow, and S3, S4 and Movie S1).

The aggregates were not surrounded by membranes, though other organelles were nearby. Vaults were seen embedded in the aggregates (Figure S3). Earlier correlative immunofluorescence/EM analyses of fixed and stained HeLa cells revealed that YFP-rhTRIM5α fluorescent bodies colocalize with the protein p62, a central component of the structure which was named sequestosome, which was also found within membrane-free compartments^43^. For this reason, we also call our aggregates sequestosomes.

Previous work showed that microtubules are required for TRIM5α motility ^11^, and we found microtubules in close association with the sequestosomes in our cryotomograms. In some cases, the microtubules were parallel to the longer dimension of the sequestosomes, as if the sequestosomes were perhaps moving along the microtubule and being shaped by collisions with other material. In many cases, the sequestosomes were also surrounded by (Figures 2 and S4), or adjacent to (Figure S5) F-actin structures. Five of these sequestosomes also appeared very close to ER tubules and mitochondria (Figures 2 and S4B, blue and orange arrows, respectively). However, given that all tomograms were recorded in the cell periphery, it is not surprising to find cortical actin, microtubules, ER and mitochondria. Indeed ~50% of our control cryotomograms contained cortical actin, microtubules, ER and mitochondria. Therefore, it could be that the sequestosomes appeared close to these structures simply by chance.

### Stage 4: Phagophore complexes

Four of the 35 YFP-rhTRIM5α fluorescent bodies imaged correlated to sequestosomes with phagophores nearby (Figures 3, 4, S6 and Movies S2 and S3). We identified phagophores in our cryotomograms as flattened vesicles near YFP-rhTRIM5α fluorescence with dimensions similar to or smaller than the sequestosomes. Only one control cryotomograms of Hela cells not expressing YFP-rhTRIM5α exhibited a similar flattened phagophore-like structure. The different morphologies and sizes of the phagophores visualized in our cryotomograms suggest they represent different stages in phagophore biogenesis. The sequestosomes in these phagophore complexes appeared just like the pre-phagophore sequestosomes in texture and size. Again, vaults were present, embedded within the sequestosome (Figure S6). We also observed putative phagophores without clear sequestosomes nearby. Strikingly in the example of Figure 4, we observed in one cryotomographic slice two separate points of contact between the phagophore and the ER membrane, both bridged by two stick-like densities ~17 nm long.

#### Hexagonal lattices near phagophores and isolated in the cytoplasm

In two cryotomograms of sequestosome:phagophore complexes, hexagonal nets with a lattice spacing of ~30 nm (center of one hexagon to center of the next) were observed in the cytoplasm close to the phagophore. The largest net covered a surface area of 500 nm^2^ and was well-ordered (Figure 4). This net appeared in close proximity not only to the sequestosome but also mitochondria, ER, and the ends of the phagophore membrane. The lattice spacing of this net and that formed by purified TRIM5α hexamers^4^ was very similar: 30 +/−2 nm *in vivo* and 33 +/−2 nm *in vitro* (Figure 5). In some cases, we also observed hexagonal lattices with a larger, ~ 35 nm lattice spacing located on the inside surface of the plasma membrane (Figure 5, S7 black circles and Movie S3). Based on their lattice spacing and location on the plasma membrane, which match the hexamers of triskelions we imaged at the plasma membrane in a previous study (Figure S5J-L)^22^, we interpret these structures as clathrin. Unlike clathrin, the smaller 30-nm nets we observed within the YFP-rhTRIM5α fluorescence signals were sandwiched between cellular material such as ribosomes and microtubules, not on the plasma membrane (Movie S3). The 30-nm nets were also not surrounding vesicles, as would be expected for intracellular clathrin, but were rather in the cytoplasm either unassociated with any membrane or found between ER tubules. Because their lattice spacing matches TRIM5α lattices *in vitro*, and they co-localized with YFP-rhTRIM5α fluorescence, we conclude they are YFP-TRIM5α.

**Figure 5.**
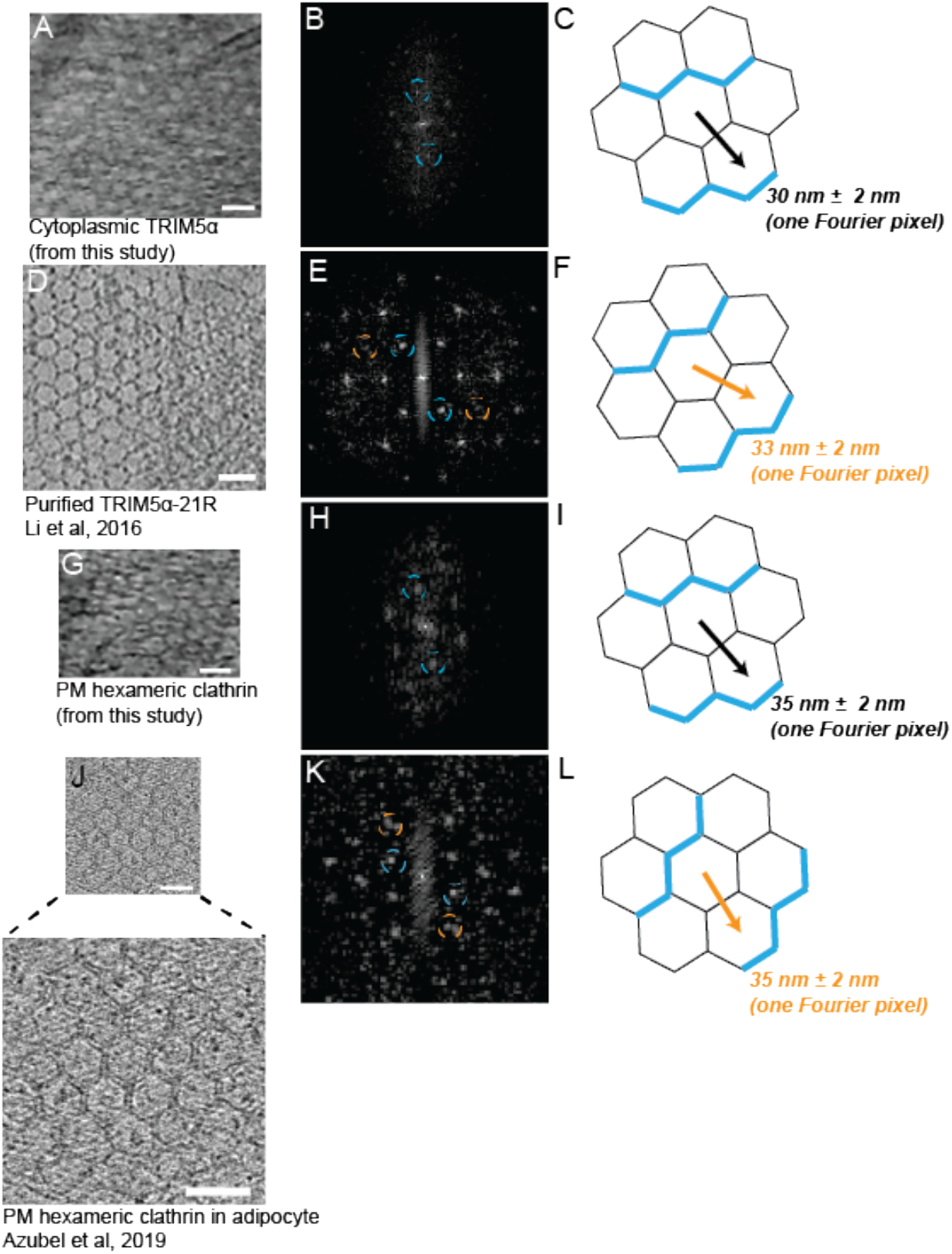
Hexagonal clathrin and TRIM5α exhibit different lattice spacing’s. **(A)** and **(D)** Cryotomographic slice showing cytoplasmic hexagonal TRIM5α inside a HeLa cell and 2D hexagonal lattices of purified TRIM5α-21R, respectively. **(G)** and **(J)** Cryotomographic slices showing hexagonal clathrin inside a HeLa and adipocyte cell respectively. **(B)**, **(E)**, **(H)**, and **(K)** Respective Fourier transfer from crytomographic slices in **(A)**, **(D)**, **(G)**, and **(J)**. Colored circles in **(B)**, **(E)**, **(H)**, and **(K)** correspond to the colored vectors and lattice spacings in **(C), (F), (I)**, and **(L)**.

### Stage 5: Autophagosomes

Seven YFP-rhTRIM5α fluorescent bodies co-localized to double-membraned vesicles (Figures 6, S8 and S9). Only one control cryotomogram exhibited a similar structure. We interpret these as autophagosomes, organelles generated by the extension and fusion of the extreme ends of the phagophore. The autophagosomal membranes displayed variable intermembrane spaces, some with large gaps devoid of macromolecular material such as ribosomes or microtubules (Figure S9). The texture of the material inside the autophagosomes was just like the cytoplasmic sequestosomes described earlier, except there were no filaments. Vaults were also seen inside the autophagosomes (Figure 6).

**Figure 6.**
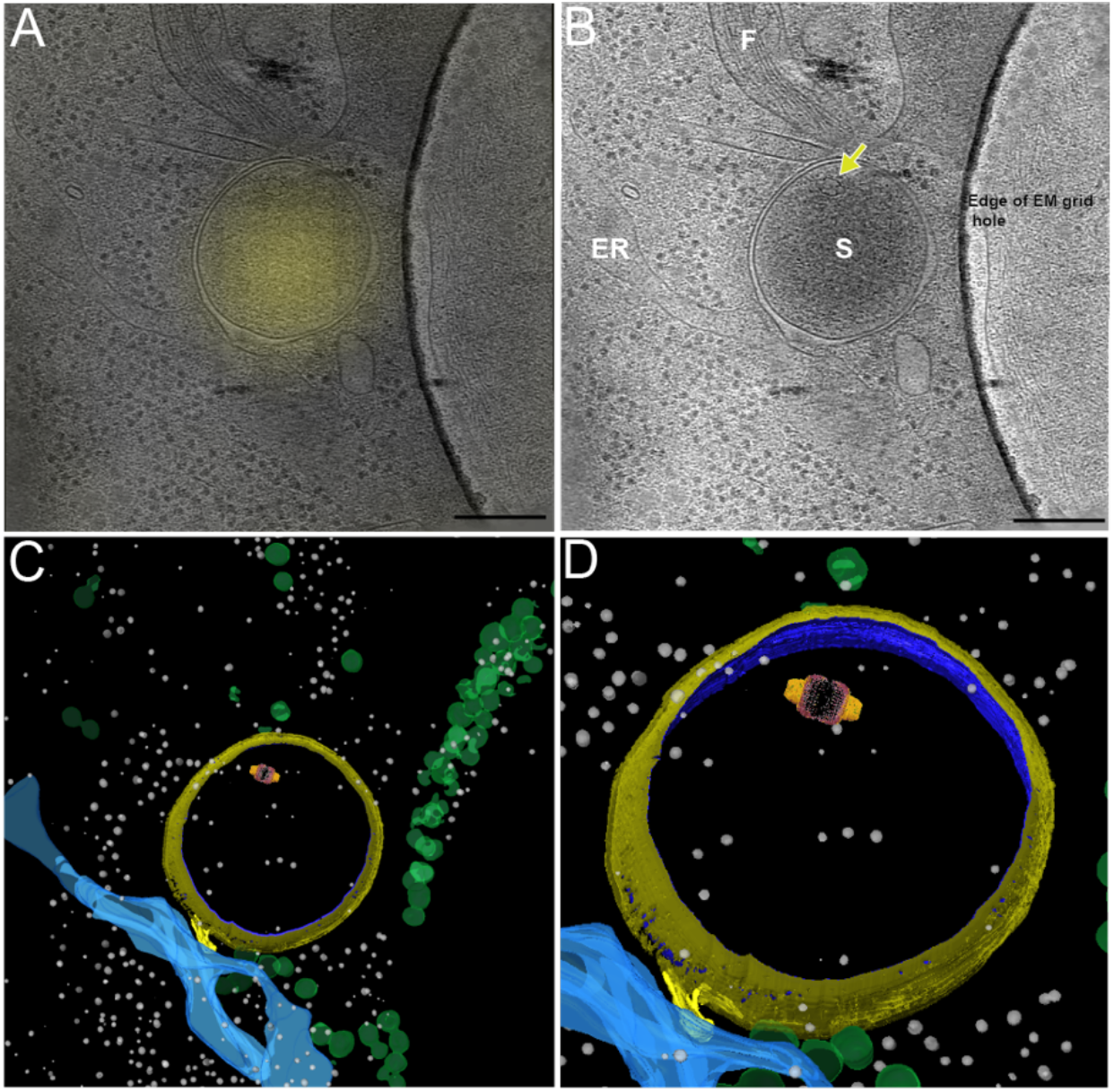
Cryo-CLEM reveals YFP-rhTRIM5α fluorescent bodies (untreated) localized to an autophagosome containing a vault. **(B)** Yellow arrow highlights a vault. Scale bars = 250 nm. **(D)** Enlarged view of the autophagosome shown in **(C)**. The crystal structure (4V60) of a vault is overlaid in **(C)** and **(D)**.

### Stage 6: Autophagic vacuoles

In two cryotomograms, we saw large (~700 nm) vesicles enclosing three or more medium-sized (~300 nm) vesicles and various cellular debris such as filaments and small (~40 nm) vesicles (Figure 7). In these two cases, the YFP-rhTRIM5α fluorescence localized specifically to one of the medium-sized vesicles. The interiors of these particular vesicles (that correlated with YFP-TRIM5α) had the same texture as the cytoplasmic sequestosomes, and they contained vaults (Figures 7 and S10). We interpret the large vesicles to be auto-lysosomes or amphisomes, and the YFP-TRIM5α-filled medium-sized vesicles to be remnants of autophagosomes after their outer membranes fused with the lysosome or early/late endosomes, respectively.

**Figure 7.**
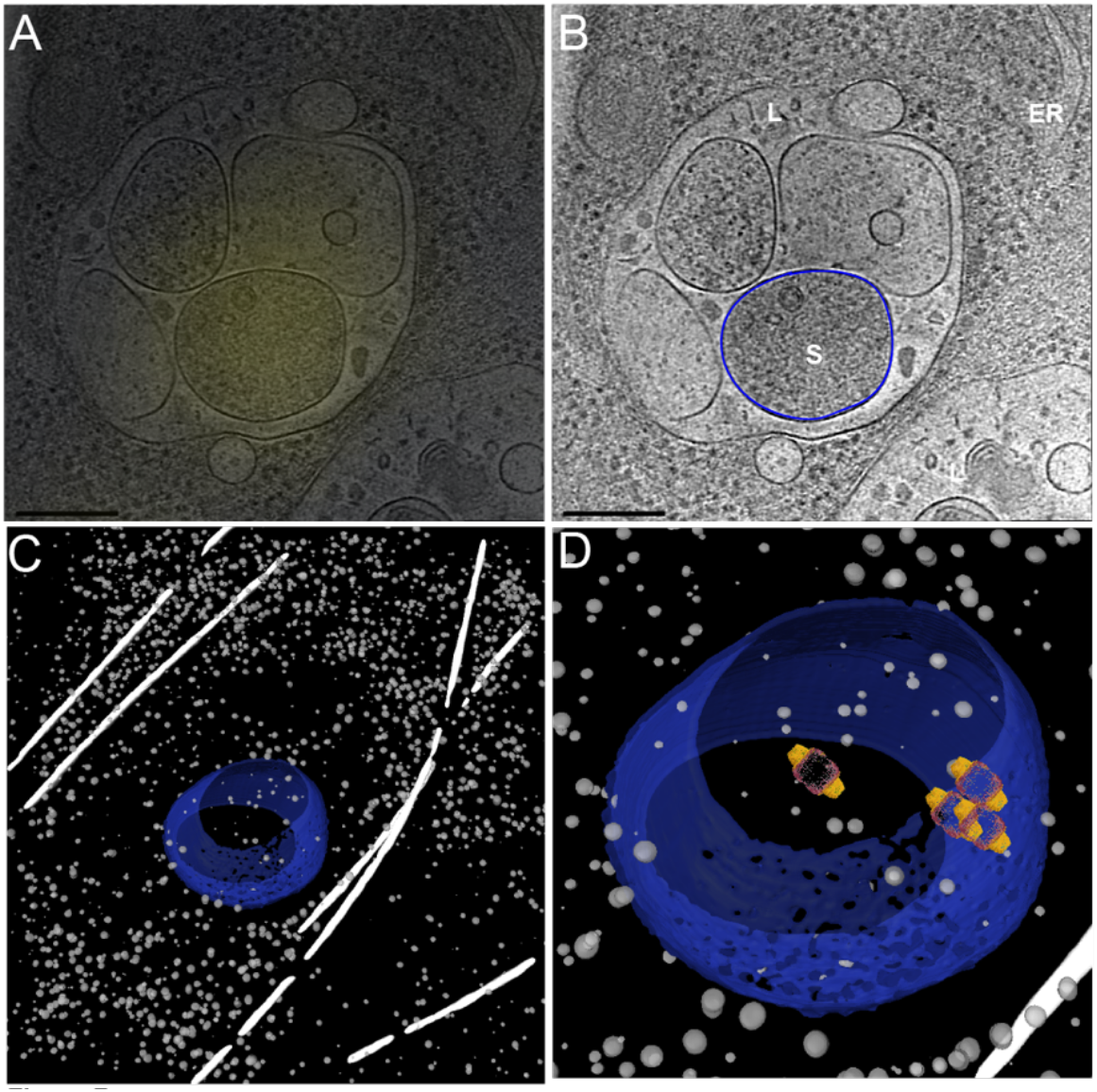
Cryo-CLEM reveals YFP-rhTRIM5α fluorescent bodies (untreated) localized to autophagic vacuoles, and autophagic vacuoles contain vaults. **(B)** The boarder of the enveloped sequestosome is highlighted with a blue line. Scale bars = 250 nm. **(D)** Enlarged view of the autophagic vacuoles outlined in blue, with the crystal structure (4V60) of the vault complex overlaid.

## Discussion

Here we found that exogenous YFP-rhTRIM5α colocalizes with six different stages of autophagic structures. We found that vaults are recruited early in autophagy, and that YFP-rhTRIM5α can form extended hexagonal nets *in vivo*.

Our results relate to several different bodies of work in the literature. First, our results support a recent report by Fletcher *et al.* that demonstrated the existence of two populations of cytoplasmic TRIM5α bodies inside cells: hexagonal signaling lattices and non-signaling aggregates^23^. We saw both extended hexagonal lattices and aggregates. Fletcher et al. showed that a single point mutation in the B-Box domain (R119E) of huTRIM5α prevented it from forming cytoplasmic fluorescent bodies or restricting retroviral infection, suggesting that pre-formed trimeric complexes within cytoplasmic fluorescent bodies are indispensable for viral restriction^23,24^. Our data support this model because here we proved by direct imaging that TRIM5α can in fact form extended hexagonal lattices inside cells.

Unfortunately, our images do not reveal what exactly the hexagonal nets are doing. Fluorescence studies have shown that overexpressed GFP-tagged human TRIM5α (GFP-huTRIM5α) localizes with DFCP1 and ULK1, two early regulators of autophagy initiation, and LC3B, a key player in phagophore biogenesis ^25,26^. A crystal structure was recently solved of the Trim5α B-box and coiled-coil regions in complex with LC3B, suggesting this contact is specific and functional ^27^. Many different receptors are used to recognize and sequester the material that is to be degraded by macroautophagy^28–30^. One of these receptors is the ubiquitin-binding p62 sequestosome protein (also known as SQSTM1). Interestingly, TRIM5α has been shown to form complexes and localize with p62 inside cells ^26,31,32^. Mandell et al. recently demonstrated that around 50 percent of TRIMs modulate autophagy by not only acting as receptors but also by forming a molecular scaffold termed the TRIMosome ^26,33,34^. Our localization of YFP-rhTRIM5α fluorescent bodies to sites of phagophore biogenesis is in agreement with all these findings, and our finding that TRIM5α can form hexagonal nets supports the notion that it may act as a scaffold. Table S1 summarizes our review of the literature which documents the association of TRIM5α proteins with many autophagy-related proteins.

Concerning the aggregates, in mammalian cells at least two kinds of aggregation compartments have been distinguished, namely the aggresome ^37,38^ and the aggresome-like induced structure (ALIS) ^39,40^. Unlike our YFP-rhTRIM5α aggregates, aggresomes persist in only one or two copies per cell at the microtubule organizing center near the nucleus and are surrounded by vimentin. ALIS have been shown to be formed by the aggregation of newly synthesized, ubiquitinated proteins, and are induced by various cell stressors including puromycin and oxidative stress ^40^. ALIS are also transient and found throughout the cell. Previous reports have shown that p62 is required for their formation ^35,40,41^ and more ALIS form when p62 is up-regulated ^42^. Indeed, ALIS are indistinguishable from p62 sequestosomes^42^. Therefore, based on these similarities, we speculate that our TRIM5α-YFP aggregates are likely ALIS rather than aggresomes. The dot-like densities in our aggregates are presumably mis-folded protein including YFP-TRIM5α ^23^.

Our data contain 3D images of putative phagophores at various stages of their growth. Strikingly, we did not see any ultrastructure in our cryotomograms like the previously described omegasome ^46^. The absence of the omegasome in our cryotomograms indicates we were either imaging a different macroautophagy pathway or the omegasome was an artifact of traditional preparative methods ^47^. Future cryo-CLEM work with tagged proteins specific for omegasome formation, such as DFCP-1 ^48^ should clarify the issue.

## Supporting information

MoviesS1

MoviesS3

## Acknowledgements

We would like to thank Sarah Speed for valuable help segmenting cryotomograms. This work was supported in part by NIH grant AI150464 to GJJ and TJH.

## Author contributions

SDC, JM, TJH and GJJ designed the experiments. SDC prepared the samples. SDC performed the cryo-CLEM. SDC, JM, TJH and GJJ interpreted the data. SDC and GJJ wrote the manuscript and the other co-authors edited and provided critical comments.

## Supplementary Materials

Figs. S1-S10

Table S1

Movies S1 – S3

---

**Figure. S1.**
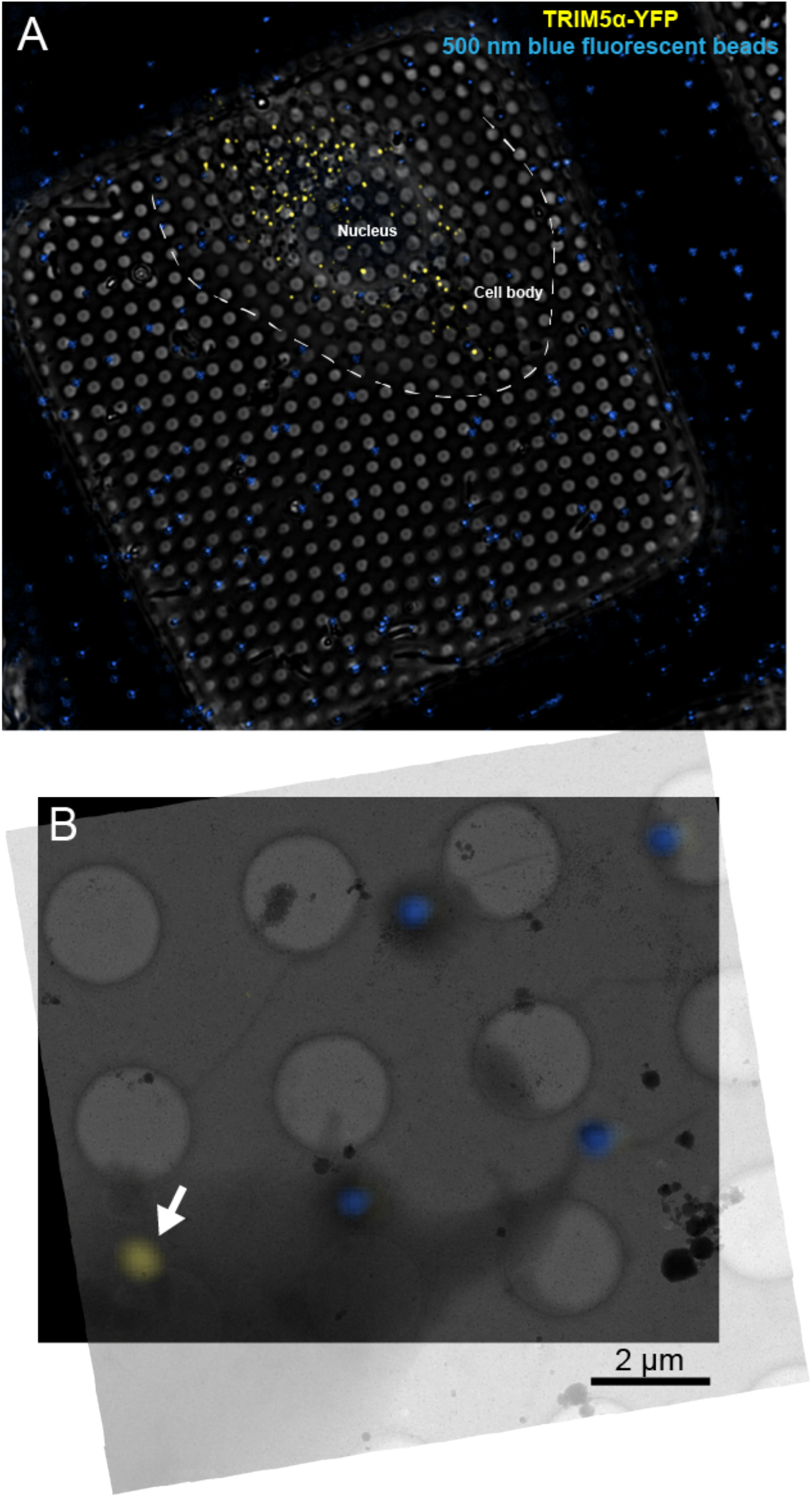
Cell line25 exhibits strong YFP-TRIM5α fluorescent bodies at 80 K. **(A)** Deconvolved cryo-LM images (composite of phase contrast, and epifluorescence in YFP and DAPI channels), of HeLa cells stably expressing YFP-TRIM5α grown on an EM grids with 500 nm blue beads added before freezing. **(B)** Bottom panel shows deconvolved epifluorescence images overlaid onto a low-magnification cryo-EM projection image. The white arrow highlights a YFP-TRIM5α fluorescent body. Scale bar = 2 μm.

**Figure. S2.**
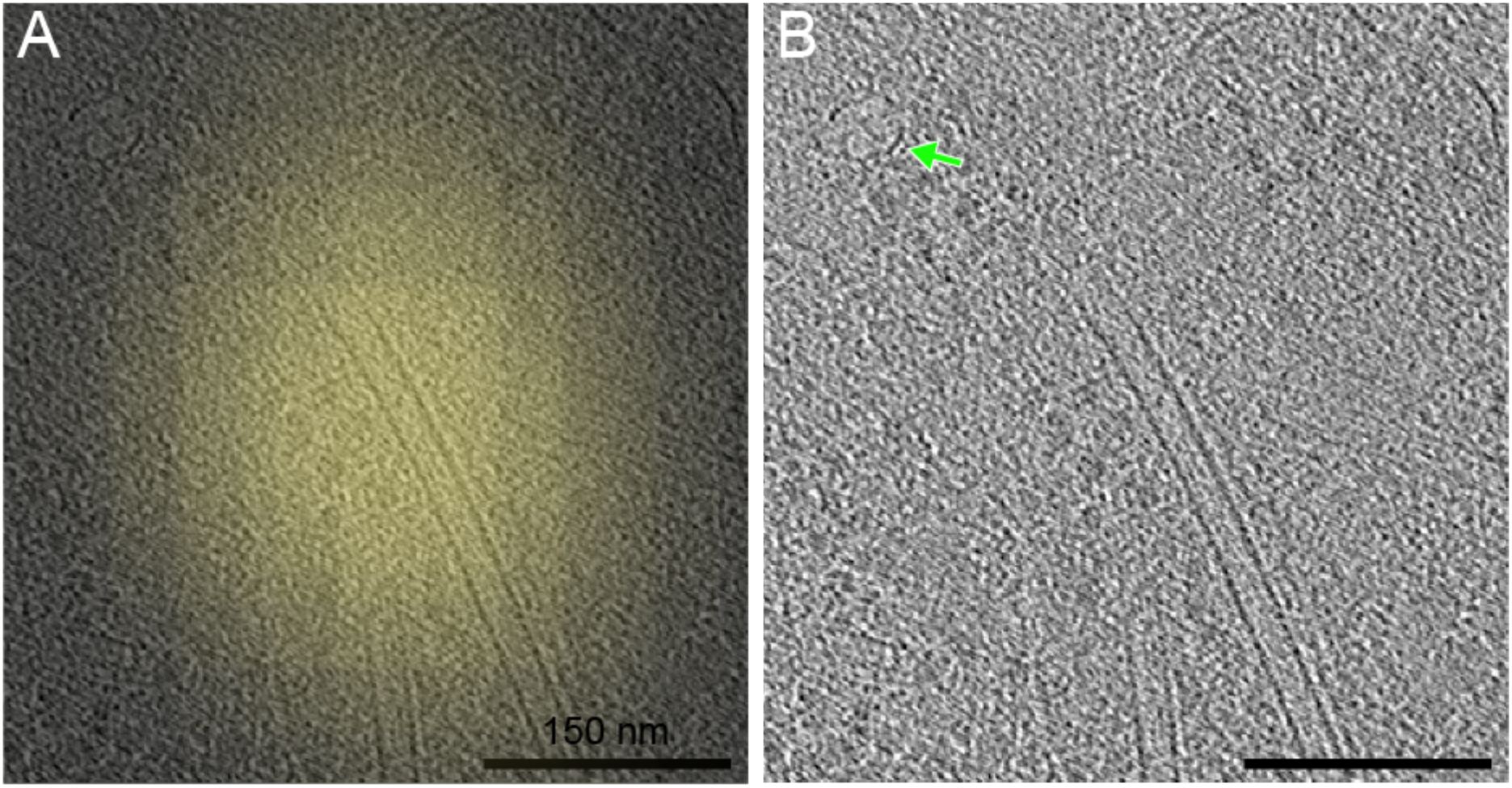
Cryo-CLEM reveals YFP-TRIM5α fluorescent bodies localize near vault complexes. The green arrow highlights a vault complex. Scale bars = 150 nm.

**Figure S3.**
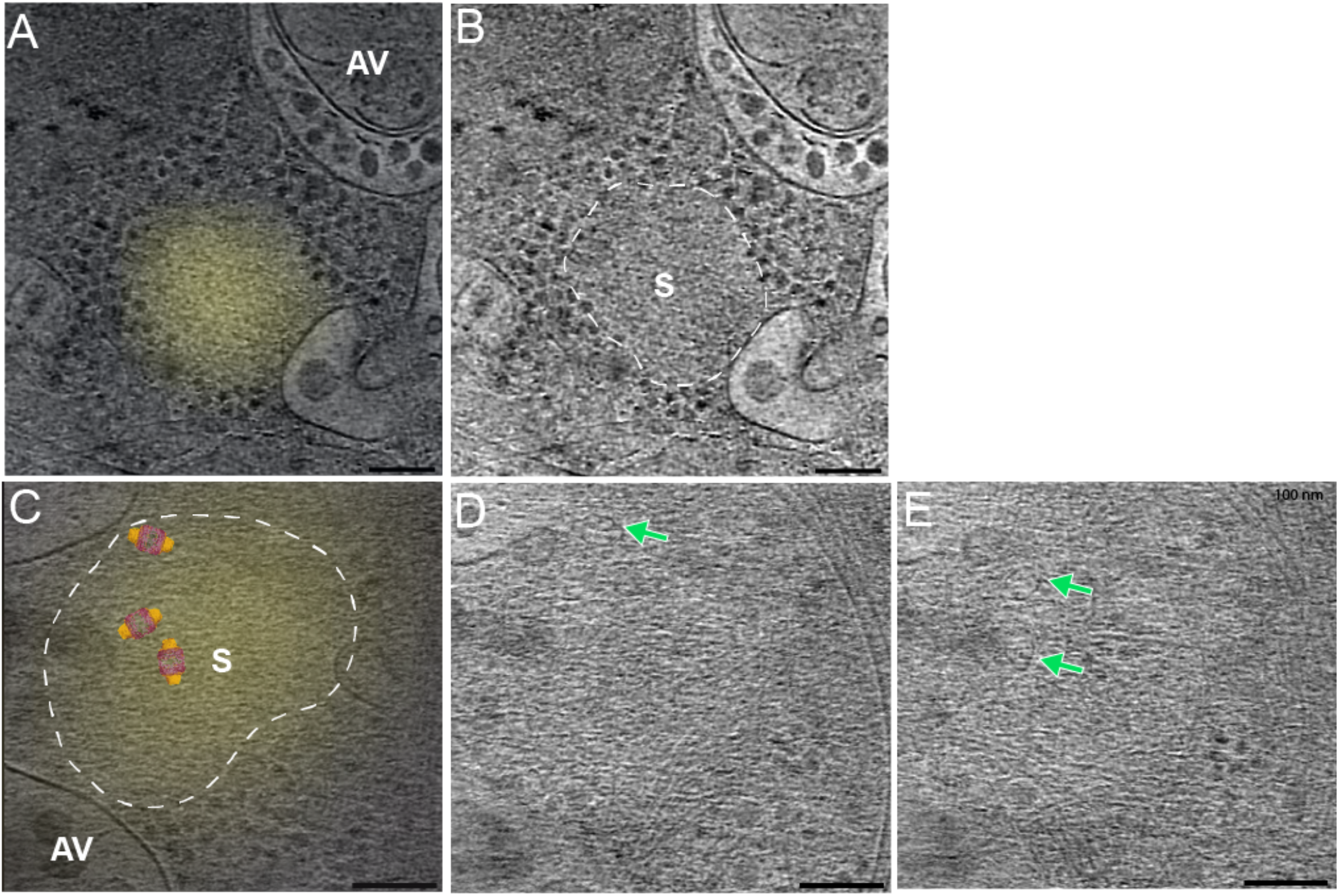
Cryo-CLEM reveals YFP-TRIM5α fluorescent bodies expressed in HeLa cells localize to cytosolic sequestosomes. **(A) and (C)** Left panel shows deconvolved epifluorescent images overlaid onto a high-magnification cryotomogram slice. **(B)** Middle panel shows the cryotomographic slice without fluorescence with the boarder of the sequestosome highlighted with a white dashed line. **(C)** The three vaults seen in **(D)** and **(E)** (different cryotomographic slices), have crystal structures (4V60) overlaid onto the cryotomogram. Cellular structures are labelled as followed S (sequestosome), and AV (autophagic vacuole). Scale bars = 100 nm. Green arrows highlight vault complexes.

**Figure S4.**
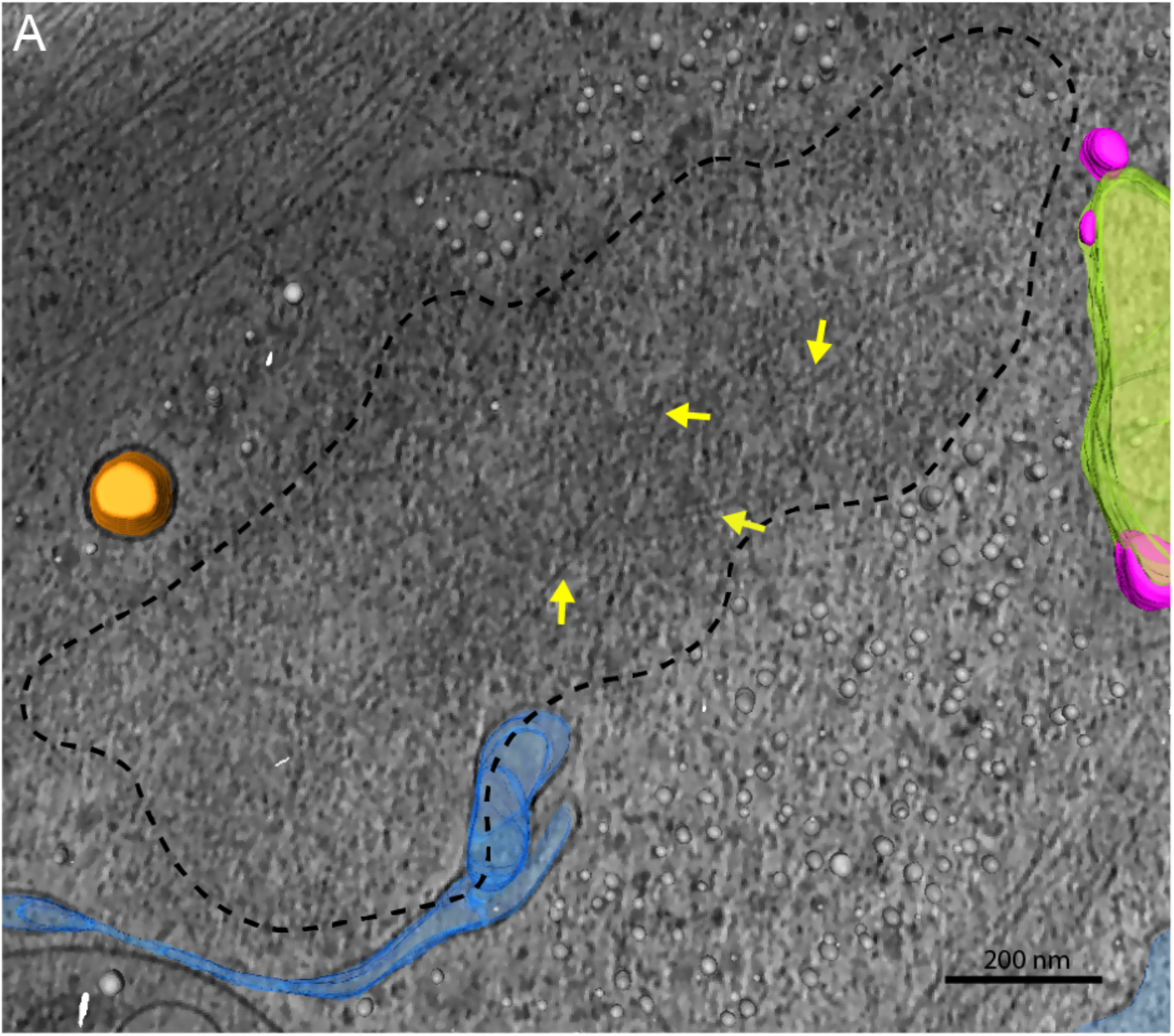
Cryo-CLEM reveals YFP-TRIM5α fluorescent bodies (MG-132), represent cytosolic sequestosomes in association with actin filaments. **(B)** The boarder of the sequestosome is highlighted with a black dashed line. Cellular structures are labelled as followed ER (endoplasmic reticulum), M (mitochondrion), and MT (microtubule). The orange arrow highlights a mitochondrion/sequestosome in close proximity, the blue arrow highlights an ER/sequestosome in close proximity, and the green arrow highlight an actin filament/sequestosome in close proximity. Scale bars = 200 nm. **(C)** Segmentation of the 3D ultrastructure inside the cell. **(D)** Bottom right panel shows a zoomed view of the area of the sequestosome shown in **(B)** and **(C)**.

**Figure S5.**
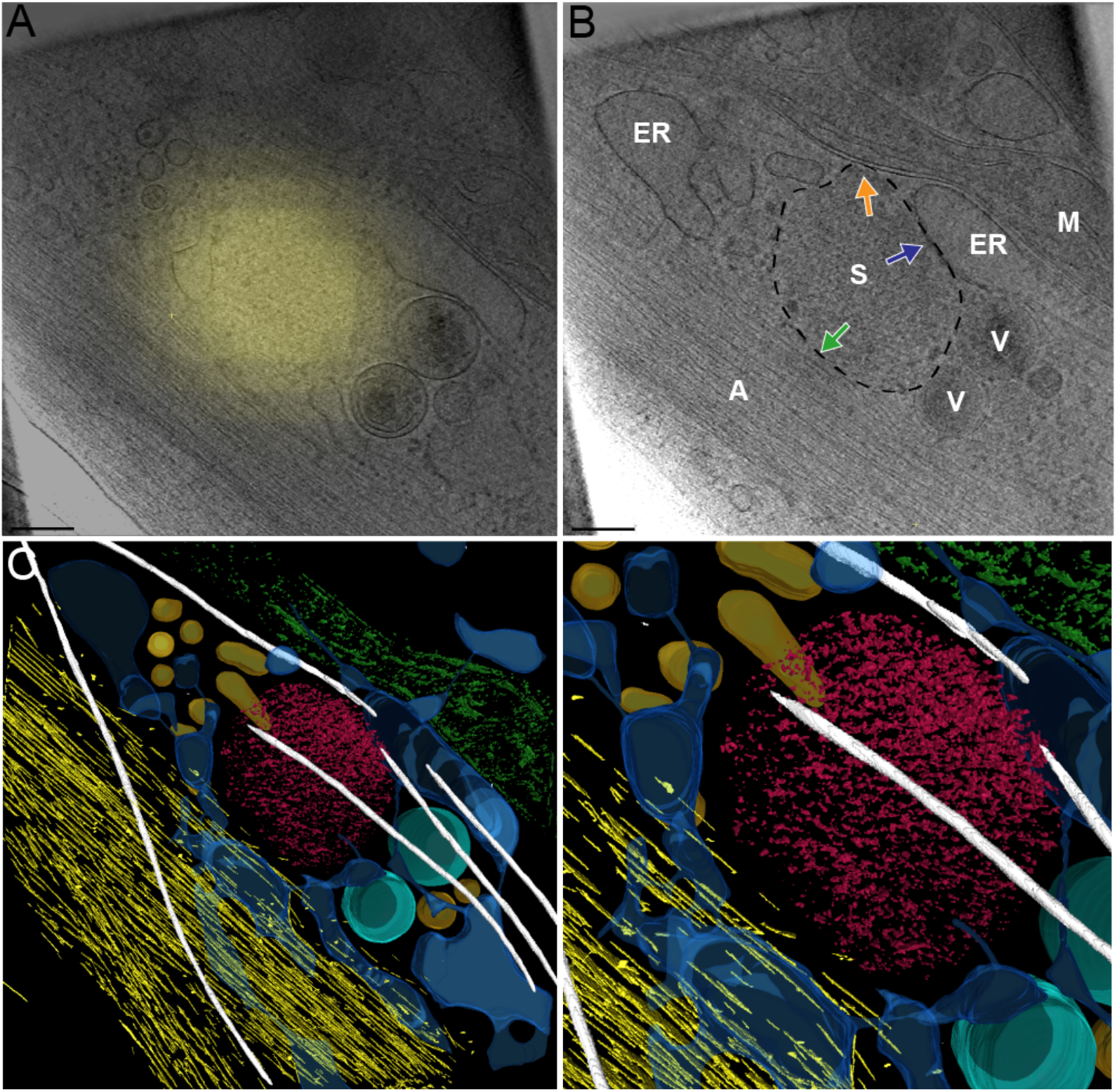
Actin filaments are observed inside sequestosomes. **(A)** A cryotomogram slice of the sequestosome outlined by a black dashed line. Yellow arrows point to areas of the sequestosome where actin filaments can be found. Scale bar = 200 nm.

**Figure S6.**
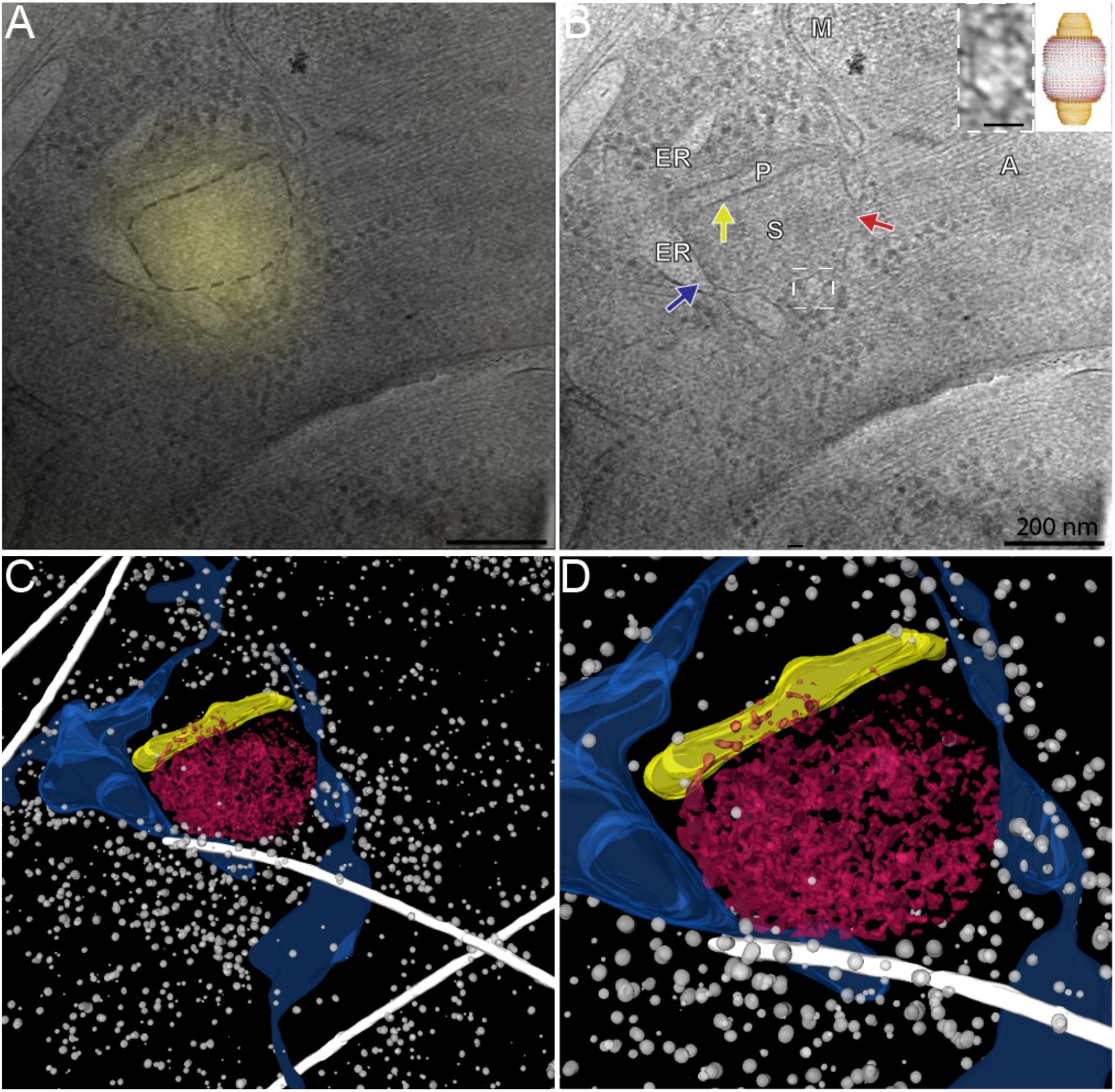
Cryo-CLEM reveals YFP-rhTRIM5α fluorescent bodies (untreated), localized to a cytosolic sequestosome:phagophore complex. **(B)** The yellow arrow highlights a sequestosome:phagophore in close proximity, the red arrow highlights a cytoplasm:sequestosome in close proximity, the blue arrow highlights a ER:sequestosome association. Scale bars = 200 nm. Inset shows an enlarged view of a vault, highlighted with a white dashed box, alongside the crystal structure (PDB:4V60) of the vault complex. Scale bar = 20 nm. **(D)** Enlarged view of the sequestosome:phagophore contact site shown in **(B)** and **(C)**.

**Figure S7.**
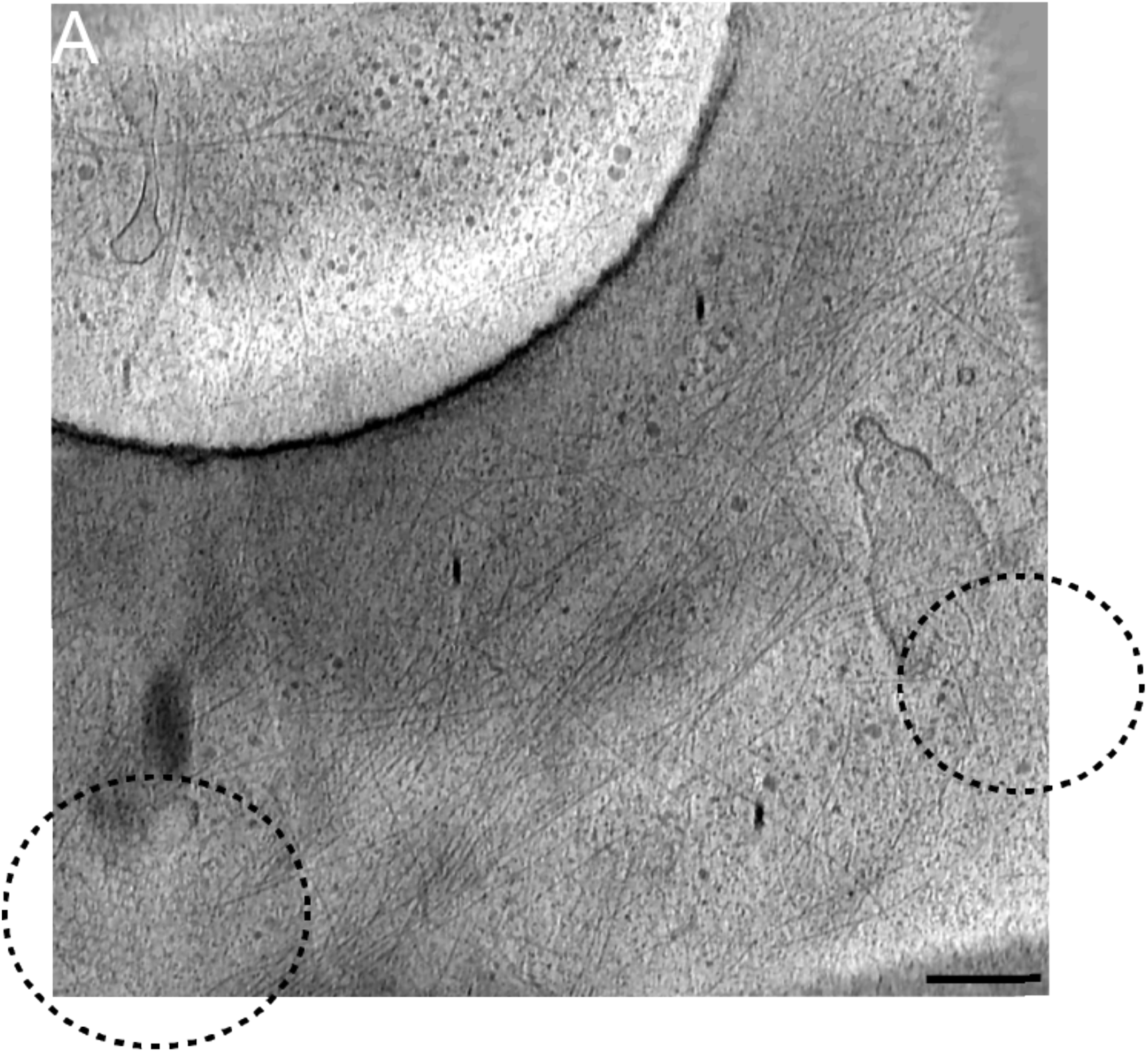
Hexagonal clathrin is located at the plasma membrane and had nearly identical architecture and dimensions to putative TRIM5α hexagonal nets found located in the cytoplasm. **(A)** Cryotomographic slice showing hexagonal clathrin at the plasma membrane circled in black. Scale bar = 250 nm.

**Figure S8.**
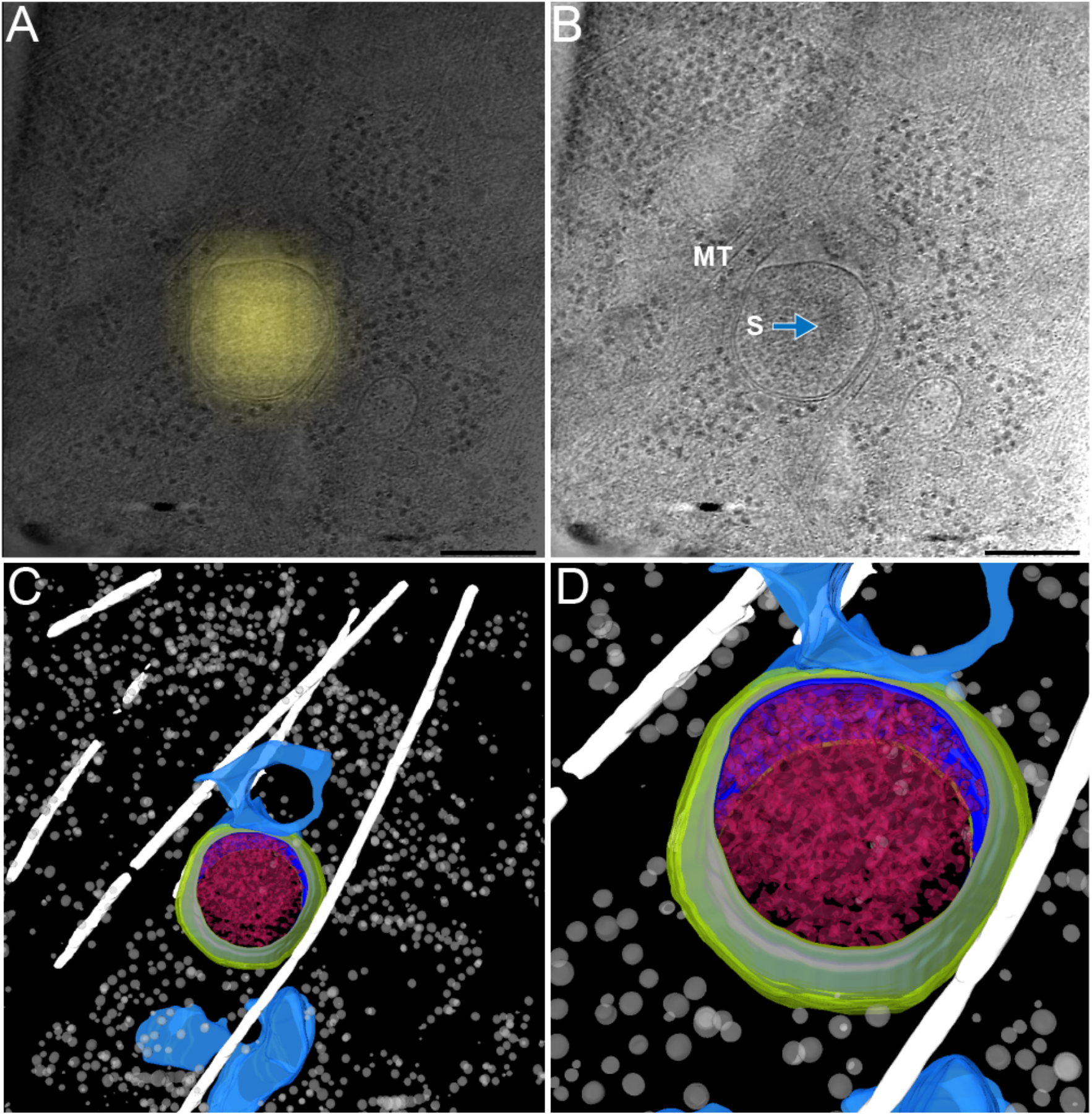
Cryo-CLEM reveals YFP-TRIM5α fluorescent bodies (untreated) localized to an autophagosome. **(A)** Upper left panel shows a deconvolved epifluorescent image overlaid onto a high-magnification cryotomogram slice. **(B)** Upper right panel shows the same cryotomographic slice without fluorescence. Blue arrow points to the sequestosome inside the autophagosome. Scale bars = 250 nm. **(C)** Bottom left panel shows an isosurface of the 3D ultrastructure inside the cell. **(D)** Bottom right panel shows a zoomed view of the enveloped sequestosome shown in **(B)** and **(C)**.

**Figure S9.**
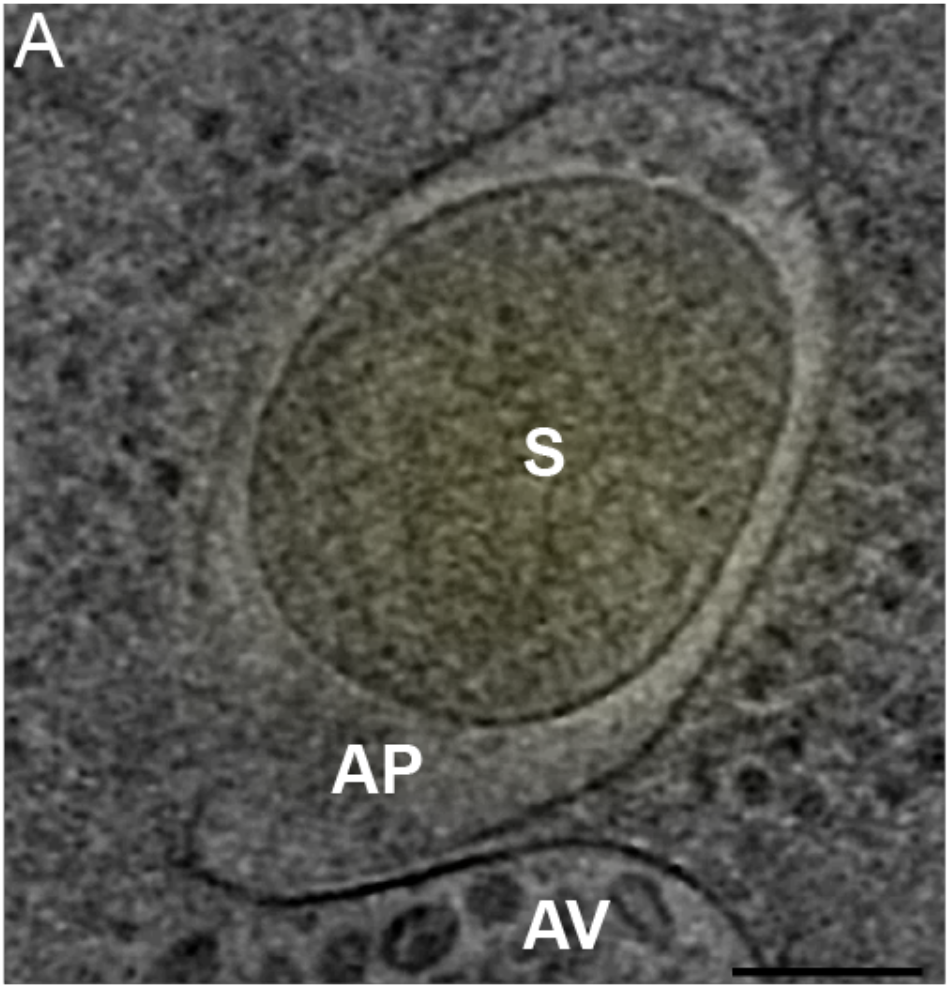
Cryo-CLEM reveals YFP-TRIM5α fluorescent bodies (untreated) localized to an autophagosome. **(A)** Upper left panel shows a deconvolved epifluorescent image overlaid onto a high-magnification cryotomogram slice. Cellular structures are labelled as followed S (sequestosome), AP (autophagosome), and AV (autophagic vacuole).

**Figure S10.**
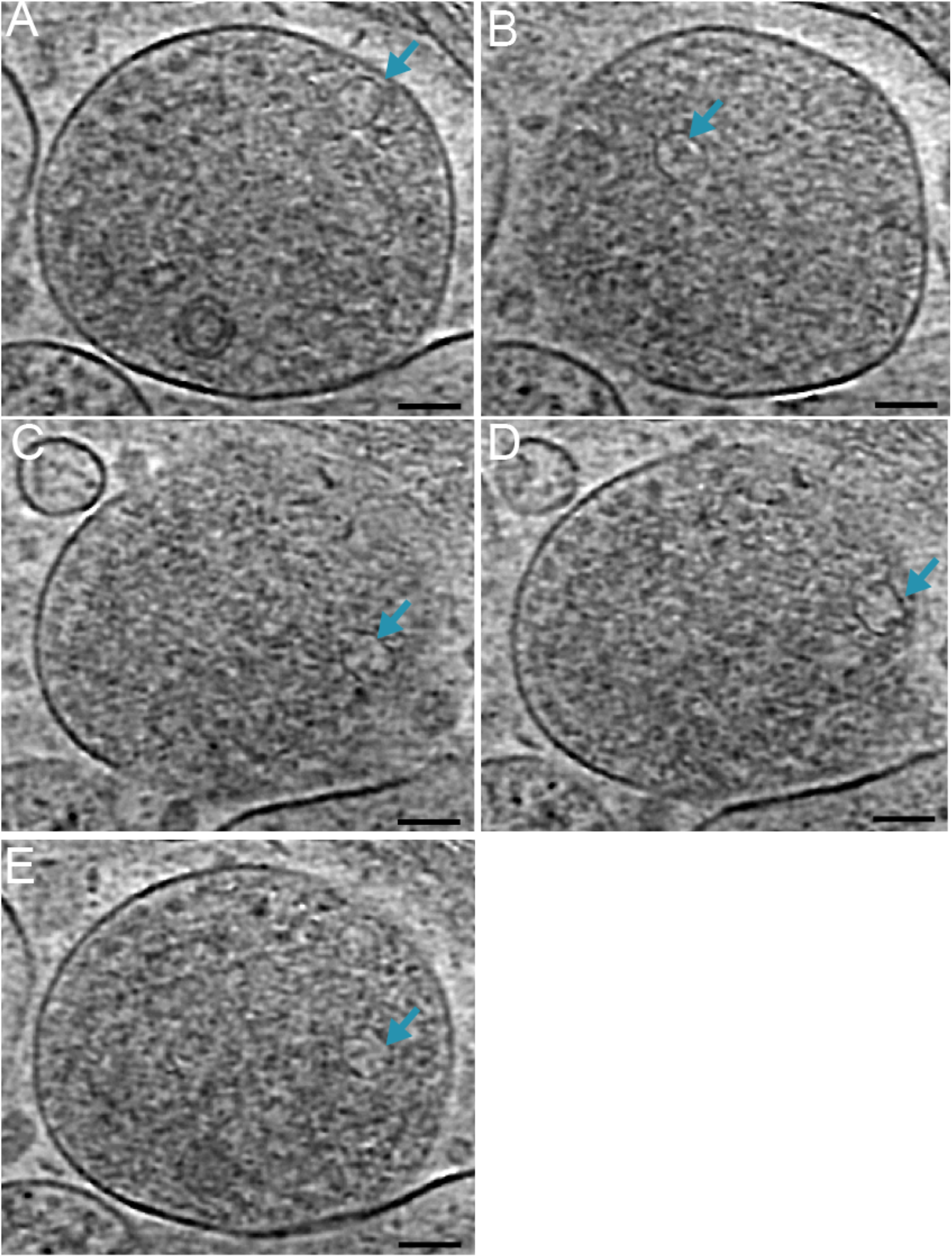
Vaults are found inside an autophagic vacuole. **(A-E)** Different tomographic slices of the autophagic vacuole in figure 7, showing 5 vaults crystal structures overlaid within the lumen of the inner portion of a post-fused autophagosome.

**Table S1.** Table detailing previous described reports of TRIM5α in proximity with other autophagy-related proteins

**Table S2.** Table detailing the cellular structures that were imaged from cells treated with MG-132 or left untreated.

## Methods

HeLa cells (cell line 25) ^12^ were maintained in a humidified 37 °C incubator with 5% CO_2_. HeLa cells were cultured where cultured in DMEM media with no phenol red (Gibco), containing 10% FBS, 100 units/mL penicillin, 100 μg/mL streptomycin. For cryo-FM and cryo-ET, cells were plated onto fibronectin-coated 200 mesh gold R2/2 London finder Quantifoil grids (Quantifoil Micro Tools GmbH, Jena, Germany) at a density of 2 × 10^5^ cells/mL. After 12 h incubation cultures were treated with MG-132 (1 μg/ml) for 2-3 hours or left untreated, before being plunge frozen in liquid ethane/propane mixture using a Vitrobot Mark IV (FEI, Hillsboro, OR) (Iancu et al., 2006). Immediately prior to plunge-freezing, 3 *μ*l of a suspension of beads was applied to grids. The bead suspension was made by diluting 500 nm blue (345/435 nm) polystyrene fluorospheres (Phosphorex) with a colloidal solution of 20 nm gold fiducials (Sigma Aldrich) pretreated with bovine serum albumin. The gold served as fiducial markers for tomogram reconstruction while the blue fluorospheres served as landmarks for registering fluorescence light microscopy (FLM) images from different channels as well as EM images ^20^. In addition, the blue fluorospheres helped locate target areas in phase contrast light microscopy and low-magnification EM images containing thin ice suitable for high-resolution ECT. Plunge-frozen grids were subsequently loaded into Polara EM cartridges (FEI). EM cartridges containing frozen grids were stored in liquid nitrogen and maintained at ≤−150 °C throughout the experiment including cryo-FLM imaging, cryo-EM imaging, storage and transfer.

### Fluorescence imaging and image processing

The EM cartridges were transferred into a cryo-FLM stage (FEI Cryostage) modified to hold Polara EM cartridges ^14,49^, and mounted on a Nikon Ti inverted microscope. The grids were imaged using a 60X extra-long-working-distance air-objective (Nikon CFI S Plan Fluor ELWD 60X NA 0.7 WD 2.62–1.8 mm). Images were recorded using a Neo 5.5 sCMOS camera (Andor Technology, South Windsor, CT) using a 2D real-time deblur deconvolution module in the NIS Elements software from AutoQuant (Nikon Instruments Inc., Melville, NY). The 2D real-time deconvolution algorithm estimates a PSF using several factors such as sample thickness, noise levels in the image, background subtraction and contrast enhancement. All fluorescence images (individual channels) were saved in 16-bit grayscale format. YFP-rhTRIM5α was visualized with a YFP filter. Blue fluorospheres were visualized with a DAPI filter

### EM imaging

Grids previously imaged by cryo-FLM were subsequently imaged by ECT using an FEI G2 Polara 300 kV FEG TEM equipped with an energy filter (slit width 20 eV for higher magnifications; Gatan, Inc.). Images were recorded using a 4 k × 4 k K2 Summit direct detector (Gatan, Inc.) operating in the electron counting mode. First, areas containing the fluorescent bodies of interest were located in the TEM using methods described previously ^20^. Tilt series were then recorded of these areas using UCSF Tomography ^50^ or SerialEM ^51^ software at a magnification of 18,000×, 27500×, or 34000×. This corresponds to a pixel size of 6 Å, 3.9 Å and 3.2 Å respectively at the specimen level and was found to be sufficient for this study. Each tilt series was collected from −60° to +60° with an increment of 1° in an automated fashion at 5–10 μm underfocus. The cumulative dose of one tilt-series was between 80 and 200 e^−^/Å^2^. The tilt series was aligned and binned by 4 into 1k x 1k using the IMOD software package ^52^, and 3D reconstructions were calculated using the simultaneous reconstruction technique (SIRT) implemented in the TOMO3D software package ^53^, or weighted back projection using IMOD. Noise reduction was performed using the non-linear anisotropic diffusion (NAD) method in IMOD ^52^, typically using a K value of 0.03–0.04 with 10 iterations.

### Segmentation and isosurface generation

Segmentation and isosurface rendering were performed in Amira (FEI). The ER, phagophores, autophagosomes, autophagic vacuoles, mitochondria and vesicles, were segmented manually using the thresholding tool. The actin filaments were segmented using the Amira XTracing Extension in Amira (FEI) ^54,55^. The microtubules and ribosomes were segmented using the tomoseg machine learning module in EMAN2 ^56,57^. Movie image sequences were generated in JPEG format in Amira (FEI) and IMOD, then converted into movies using QuickTime Player 7. Photoshop CS6 (Adobe) was then used to produce the final versions of the movies.

## Supplementary figures and table

**Table S1.** TRIM5α interactions inside cells

| Cell Type and Method | Interaction | Paper |
| --- | --- | --- |
| 293T<br>Immunoprecipitation assay | TAK1, TAB2 and TAB3 | Pertel et al. 2011 |
| HeLa<br>Fluorescence microscopy<br>and Immunoprecipitation<br>assay | p62 | Mandell et al. 2014,<br>O'Connor et al. 2010 |
| Fluorescence microscopy | TRIM17 | Mandell et al. 2014,<br>Mandell et al. 2016 |
| Coimmunoprecipitation<br>X-ray crystallography<br>Fluorescence microscopy | ATG8 paralogs -<br>GABARAP GABARAPL1<br>LC3A, LC3B, LC3C and<br>GABARAPL2 | Mandell et al. 2014<br>Keown et al. 2018 |
| HeLa cells<br>Fluorescence microscopy | DFCP1 omegasome | Mandell et al. 2014 |
| HeLa cells Fluorescence<br>microscopy and 293T<br>Coimmunoprecipitation<br>assay | ULK1 | Mandell et al. 2014 |
| Proximity ligation assay | Beclin 1 | Mandell et al. 2014 |
| Immunoprecipitation assay | p62 | Ribero et al. 2016 |
| Immunoprecipitation assay | Atg16L1 | Ribero et al. 2016 |
| Immunoprecipitation assay | Atg5 | Ribero et al. 2016 |

**Table S2.** Cellular structures and autophagic organelles found with and without MG-132

| Cell | MG-132 Treatment | Cellular structure |
| --- | --- | --- |
| 1 | - | Sequestosome |
| 1 | - | 2 x Autophagosome |
| 2 | - | Sequestosome |
| 2 | - | Phagophore |
| 3 | - | Autophagosome |
| 3 | - | Autophagic Vacuoles |
| 3 | - | Phagophore/Sequestosome |
| 4 | - | Autophagosome |
| 4 | - | Autophagosome |
| 5 | - | TRIM5α hexagons |
| 6 | - | Phagophore/Sequestosome |
| 6 | - | 2 x Sequestosome |
| 7 | - | Autophagic Vacuoles |
| 7 |  | Vault |
| 8 | - | Autophagosome |
| 9 | + | 2 x Autophagic Vacuoles |
| 9 | + | Phagophore/Hexagonal Net |
| 10 | + | Phagophore/Sequestosome |
| 11 | + | Sequestosome |
| 12 | + | Autophagic Vacuoles |
| 13 | + | Sequestosome/Vaults |
| 14 | + | Sequestosome |
| 15 | + | Sequestosome |
| 16 | + | Sequestosome |
| 17 | + | Phagophore/Sequestosome |
| 18 | + | Sequestosome |
| 19 | + | Autophagosome |
| 20 | + | Vault |

## References

1. Reymond, A. et al. The tripartite motif family identifies cell compartments. EMBO J. 20, 2140–2151 (2001).

2. Stremlau, M. et al. The cytoplasmic body component TRIM5alpha restricts HIV-1 infection in Old World monkeys. Nature 427, 848–853 (2004).

3. Ganser-Pornillos, B. K. et al. Hexagonal assembly of a restricting TRIM5α protein. Proc. Natl. Acad. Sci. 108, 534–539 (2011).

4. Li, Y.-L. et al. Primate TRIM5 proteins form hexagonal nets on HIV-1 capsids. eLife 5, e16269 (2016).

5. Sanchez, J. G. et al. The tripartite motif coiled-coil is an elongated antiparallel hairpin dimer. Proc. Natl. Acad. Sci. U. S. A. 111, 2494–2499 (2014).

6. Wagner, J. M. et al. Mechanism of B-box 2 domain-mediated higher-order assembly of the retroviral restriction factor TRIM5α. eLife 5, (2016).

7. Nisole, S., Lynch, C., Stoye, J. P. & Yap, M. W. A Trim5-cyclophilin A fusion protein found in owl monkey kidney cells can restrict HIV-1. Proc. Natl. Acad. Sci. U. S. A. 101, 13324–13328 (2004).

8. Sayah, D. M., Sokolskaja, E., Berthoux, L. & Luban, J. Cyclophilin A retrotransposition into TRIM5 explains owl monkey resistance to HIV-1. Nature 430, 569–573 (2004).

9. Stremlau, M., Perron, M., Welikala, S. & Sodroski, J. Species-specific variation in the B30.2(SPRY) domain of TRIM5alpha determines the potency of human immunodeficiency virus restriction. J. Virol. 79, 3139–3145 (2005).

10. Stremlau, M. et al. Specific recognition and accelerated uncoating of retroviral capsids by the TRIM5alpha restriction factor. Proc. Natl. Acad. Sci. U. S. A. 103, 5514–5519 (2006).

11. Campbell, E. M. et al. TRIM5 alpha cytoplasmic bodies are highly dynamic structures. Mol. Biol. Cell 18, 2102–2111 (2007).

12. Campbell, E. M., Perez, O., Anderson, J. L. & Hope, T. J. Visualization of a proteasome-independent intermediate during restriction of HIV-1 by rhesus TRIM5α. J. Cell Biol. 180, 549–561 (2008).

13. Narayan, K. et al. Multi-resolution Correlative Focused Ion Beam Scanning Electron Microscopy: Applications to Cell Biology. J. Struct. Biol. 185, 278–284 (2014).

14. Briegel, A. et al. Chapter Thirteen - Correlated Light and Electron Cryo-Microscopy. in Methods in Enzymology (ed. Jensen, G. J.) 481, 317–341 (Academic Press, 2010).

15. Bjørkøy, G. et al. p62/SQSTM1 forms protein aggregates degraded by autophagy and has a protective effect on huntingtin-induced cell death. J. Cell Biol. 171, 603–614 (2005).

16. Rogov, V., Dötsch, V., Johansen, T. & Kirkin, V. Interactions between Autophagy Receptors and Ubiquitin-like Proteins Form the Molecular Basis for Selective Autophagy. Mol. Cell 53, 167–178 (2014).

17. Stolz, A., Ernst, A. & Dikic, I. Cargo recognition and trafficking in selective autophagy. Nat. Cell Biol. 16, 495–501 (2014).

18. Woodward, C. L., Mendonça, L. M. & Jensen, G. J. Direct visualization of vaults within intact cells by electron cryo-tomography. Cell. Mol. Life Sci. CMLS 72, 3401–3409 (2015).

19. Horos, R. et al. The Small Non-coding Vault RNA1-1 Acts as a Riboregulator of Autophagy. Cell 176, 1054–1067.e12 (2019).

20. Iancu, C. V. et al. Electron cryotomography sample preparation using the Vitrobot. Nat. Protoc. 1, 2813–2819 (2006).

21. Wu, X., Anderson, J. L., Campbell, E. M., Joseph, A. M. & Hope, T. J. Proteasome inhibitors uncouple rhesus TRIM5α restriction of HIV-1 reverse transcription and infection. Proc. Natl. Acad. Sci. 103, 7465–7470 (2006).

22. Carter, S. D. et al. Distinguishing signal from autofluorescence in cryogenic correlated light and electron microscopy of mammalian cells. J. Struct. Biol. 201, 15–25 (2018).

23. EMAN2: an extensible image processing suite for electron microscopy. - PubMed - NCBI. Available at: https://www.ncbi.nlm.nih.gov/pubmed/16859925. (Accessed: 6th May 2019)

24. Bjørkøy, G. et al. p62/SQSTM1 forms protein aggregates degraded by autophagy and has a protective effect on huntingtin-induced cell death. J. Cell Biol. 171, 603–614 (2005).

25. Azubel, M. et al. FGF21 trafficking in intact human cells revealed by cryo-electron tomography with gold nanoparticles. eLife 8, e43146 (2019).

26. Fletcher, A. J. et al. Trivalent RING Assembly on Retroviral Capsids Activates TRIM5 Ubiquitination and Innate Immune Signaling. Cell Host Microbe 24, 761–775.e6 (2018).

27. Diaz-Griffero, F. et al. Modulation of Retroviral Restriction and Proteasome Inhibitor-Resistant Turnover by Changes in the TRIM5α B-Box 2 Domain. J. Virol. 81, 10362–10378 (2007).

28. Itakura, E. & Mizushima, N. Characterization of autophagosome formation site by a hierarchical analysis of mammalian Atg proteins. Autophagy 6, 764–776 (2010).

29. Mandell, M. A. et al. TRIM proteins regulate autophagy and can target autophagic substrates by direct recognition. Dev. Cell 30, 394–409 (2014).

30. Keown, J. R. et al. A helical LIR mediates the interaction between the retroviral restriction factor Trim5α and the mammalian autophagy related ATG8 proteins. J. Biol. Chem. jbc.RA118.004202 (2018). doi:10.1074/jbc.RA118.004202

31. Kaur, J. & Debnath, J. Autophagy at the crossroads of catabolism and anabolism. Nat. Rev. Mol. Cell Biol. 16, 461–472 (2015).

32. Seibenhener, M. L. et al. Sequestosome 1/p62 is a polyubiquitin chain binding protein involved in ubiquitin proteasome degradation. Mol. Cell. Biol. 24, 8055–8068 (2004).

33. Wani, W. et al. Regulation of autophagy by protein post-translational modification. Lab. Investig. J. Tech. Methods Pathol. 95, 14–25 (2015).

34. O’Connor, C. et al. p62/Sequestosome-1 Associates with and Sustains the Expression of Retroviral Restriction Factor TRIM5α. J. Virol. 84, 5997–6006 (2010).

35. Ribeiro, C. M. S. et al. Receptor usage dictates HIV-1 restriction by human TRIM5α in dendritic cell subsets. Nature 540, 448–452 (2016).

36. Mandell, M. A., Kimura, T., Jain, A., Johansen, T. & Deretic, V. TRIM proteins regulate autophagy: TRIM5 is a selective autophagy receptor mediating HIV-1 restriction. Autophagy 10, 2387–2388 (2015).

37. Mandell, M. A. et al. TRIM17 contributes to autophagy of midbodies while actively sparing other targets from degradation. J. Cell Sci. 129, 3562–3573 (2016).

38. García-Mata, R., Bebök, Z., Sorscher, E. J. & Sztul, E. S. Characterization and Dynamics of Aggresome Formation by a Cytosolic Gfp-Chimera✪. J. Cell Biol. 146, 1239–1254 (1999).

39. Johnston, J. A., Ward, C. L. & Kopito, R. R. Aggresomes: a cellular response to misfolded proteins. J. Cell Biol. 143, 1883–1898 (1998).

40. Lelouard, H. et al. Transient aggregation of ubiquitinated proteins during dendritic cell maturation. Nature 417, 177–182 (2002).

41. Szeto, J. et al. ALIS are Stress-Induced Protein Storage Compartments for Substrates of the Proteasome and Autophagy. Autophagy 2, 189–199 (2006).

42. Clausen, T. H. et al. p62/SQSTM1 and ALFY interact to facilitate the formation of p62 bodies/ALIS and their degradation by autophagy. Autophagy 6, 330–344 (2010).

43. Pankiv, S. et al. p62/SQSTM1 Binds Directly to Atg8/LC3 to Facilitate Degradation of Ubiquitinated Protein Aggregates by Autophagy. J. Biol. Chem. 282, 24131–24145 (2007).

44. Liu, X.-D. et al. Transient Aggregation of Ubiquitinated Proteins Is a Cytosolic Unfolded Protein Response to Inflammation and Endoplasmic Reticulum Stress. J. Biol. Chem. 287, 19687–19698 (2012).

45. Hayashi-Nishino, M. et al. A subdomain of the endoplasmic reticulum forms a cradle for autophagosome formation. Nat. Cell Biol. 11, 1433–1437 (2009).

46. Pilhofer, M., Ladinsky, M. S., McDowall, A. W. & Jensen, G. J. Chapter 2 - Bacterial TEM: New Insights from Cryo-Microscopy. in Methods in Cell Biology (ed. Müller-Reichert, T.) 96, 21–45 (Academic Press, 2010).

47. Axe, E. L. et al. Autophagosome formation from membrane compartments enriched in phosphatidylinositol 3-phosphate and dynamically connected to the endoplasmic reticulum. J. Cell Biol. 182, 685–701 (2008).

48. Nickell, S., Kofler, C., Leis, A. P. & Baumeister, W. A visual approach to proteomics. Nat. Rev. Mol. Cell Biol. 7, 225–230 (2006).

49. Zheng, S. Q. et al. UCSF tomography: An integrated software suite for real-time electron microscopic tomographic data collection, alignment, and reconstruction. J. Struct. Biol. 157, 138–147 (2007).

50. Mastronarde, D. N. Automated electron microscope tomography using robust prediction of specimen movements. J. Struct. Biol. 152, 36–51 (2005).

51. Kremer, J. R., Mastronarde, D. N. & McIntosh, J. R. Computer visualization of three-dimensional image data using IMOD. J. Struct. Biol. 116, 71–76 (1996).

52. Agulleiro, J. I. & Fernandez, J. J. Fast tomographic reconstruction on multicore computers. Bioinforma. Oxf. Engl. 27, 582–583 (2011).

53. Rigort, A. et al. Automated segmentation of electron tomograms for a quantitative description of actin filament networks. J. Struct. Biol. 177, 135–144 (2012).

54. Weber, B. et al. Automated tracing of microtubules in electron tomograms of plastic embedded samples of Caenorhabditis elegans embryos. J. Struct. Biol. 178, 129–138 (2012).

55. Chen, M. et al. Convolutional neural networks for automated annotation of cellular cryo-electron tomograms. Nat. Methods 14, 983–985 (2017).

56. Chen, M. et al. Protocol for Convolutional Neural Networks based Automated Cellular Cryo-Electron Tomograms Annotation. Protoc. Exch. (2017). doi:10.1038/protex.2017.095

